# Integrated histopathologic modeling of detailed tumor subtypes and actionable biomarkers

**DOI:** 10.1101/2025.08.14.670351

**Authors:** Kevin M Boehm, Madison Darmofal, Arfath Pasha, Andrew Aukerman, Raymond Lim, Evan Seffar, Tom Pollard, Natasha Rekhtman, Jason Chang, Armaan Kohli, Darin Moore, Marta Ligero, JianJiong Gao, Georgios Asimomitis, Anika Begum, Fresia Pareja, Hikmat Al-Ahmadie, Klaus J Busam, Nikolaos M Dimitriou, Meera Hameed, Ahmet Dogan, Lora H Ellenson, Jie-Fu Chen, Daniel Gomez, Nancy Lee, Eric Sherman, Himanshu Nagar, Lior Z Braunstein, Britta Weigelt, James Fagin, Susan Fu, Jonathan Alarcon, Neelraj Patil, Areej Alsaafin, A. Rose Brannon, Kofi Amoah, Jordan Eichholz, Martin Voss, Justin Jee, Christopher Fong, Michele Waters, Luke R.G. Pike, Pedram Razavi, Paul Romesser, Atif Khan, Walid K Chatila, Aasiya Islam, Elana Sverdlik, Ino de Bruijn, Zain-Ul-Abideen Nasir, Karl Pichotta, Jinru Shia, Cristina R. Antonescu, Victor Reuter, Jake June-Koo Lee, Marc Ladanyi, Orly Ardon, Kojo Elenitoba-Johnson, Michael F Berger, David Solit, Nikolaus Schultz, Sohrab P Shah, Francisco Sánchez-Vega

**Author notes:** These authors contributed equally (Kevin M Boehm, Madison Darmofal, Arfath Pasha). These authors jointly supervised the work (Nikolaus Schultz, Sohrab P Shah, Francisco Sánchez-Vega).

## Abstract

Accurate cancer subtyping with accompanying molecular characterization is critical for precision oncology. While machine learning approaches have been applied to both digital pathology and cancer genomics, previous work has been limited in sample size and has typically aggregated granular cancer subtypes into coarse groupings, likely obfuscating informative molecular and prognostic associations and phenotypic variation of more detailed tumor subtypes. Accordingly, we collated 378,123 hematoxylin and eosin (H&E)-stained whole-slide images (WSIs) with matched targeted DNA clinical sequencing results and OncoTree detailed cancer subtypes from a real-world cohort of 71,142 patients. Using this scaled, granular dataset and a cancer subtype knowledge graph, we developed Mosaic: a family of calibrated machine learning models using H&E WSI embeddings to classify tumors and identify molecular phenotypes across 163 detailed subtypes. The cancer subtyping module (Aeon) achieved an area under the receiver operating characteristic curve (AUROC) of 0.992 overall, with 161/163 subtypes reaching an AUROC ≥ 0.90 and improved performance over a state-of-the-art genomics-based classifier. The genomic inference module (Paladin) achieved an AUROC ≥ 0.80 for 167 pairs of detailed subtypes and genomic targets. We further used the learned histopathologic representations to i) identify key associations of the histopathologic embeddings with clinical biomarkers; ii) identify unsupervised sub-clusters of tumors with genomic determinants of tumor phenotype; iii) specify granular diagnoses for cancers of unknown primary, evaluated by genomic associations and expected clinical outcome distributions; iv) annotate functional significance for variants of uncertain significance (VUS); and v) identify cases that mimic the phenotypic effect of known DNA variants on H&E in the absence of detectable DNA alterations. Taken together, this work advances our understanding of phenotypic variation of granular tumor subtypes, their relevance to enhanced diagnostics, and their potential utility in risk stratification with multimodal machine learning in cancer.

## Introduction

Cancer subtypes are determined by anatomic site of origin, cell of origin, and—increasingly— characteristic molecular features. Correctly diagnosing cancer subtypes is a central component of oncology that informs therapy selection and patient prognosis, motivating the development of tumor classification ontologies, such as OncoTree^1^. At the extreme end of the spectrum, cancers of unknown primary (CUP) present a diagnostic dilemma and portend poor prognosis^2–5^. This has motivated the development of molecular classifiers such as Genome-Derived-Diagnosis Ensemble (GDD-ENS) and transcriptomic assays for cancer subtype assignment^6–9^. In addition to genomic techniques, histopathology-based classifiers such as TOAD^10^ and CHIEF^11^ have been developed to infer cancer subtypes ^12–14^ from H&E WSIs. However, these methods identify only eighteen coarsely defined subtypes out of hundreds used in practice, resulting in over-aggregation of disparate histologies by anatomic site without regard for cell of origin. We posit that this approach lacks required specificity and furthermore has the potential to miss key molecular subtypes with prognostic or predictive relevance^15–17^. In this work, we therefore addressed this significant unmet need by developing more detailed, granular subtype models based on histopathological images and associated phenotypic features with genomic and other clinical covariates.

Genomic features of individual tumors add additional information on top of granular subtypes, indicating specific targeted therapy and/or prognostic risk stratification. In rare cases, genomic features are causal determinants of characteristic histopathologic appearance, yielding established phenotypes within granular cancer subtypes. For example, microsatellite instability (MSI) status in colorectal adenocarcinoma is associated with human-detectable characteristics of the tumor, including medullary features, the presence of intraepithelial lymphocytes, and poor differentiation^18^. This association motivated foundational work in digital pathology^19^ leading to European Union-approved clinical-grade inference^20–22^ of MSI, an FDA-recognised indicator of response to pembrolizumab, among other therapeutic and prognostic implications^23,24^. Additional studies have tested associations between tumor phenotypes on H&E WSIs with individual genomic biomarkers in individual cancer types^25,26^ and pan-cancer^27–30^ settings, finding that pathognomonic features of individual variants are uncommon. However, these studies were limited by their reliance upon small datasets for training and validation (most commonly using approximately 5,000 patients from The Cancer Genome Atlas, TCGA). Furthermore, studies have aggregated tumor subtypes with different prior probabilities of genomic features, which leads to models that recapitulate known phenotype-genotype associations rather than discovering new ones^11^. For example, identifying *TP53* oncogenic variants in all “ovarian cancer” amounts to identifying high-grade serous histology, which is knowable from the diagnosis alone^31^. We suggest that stratification by granular subtype will allow models to discover variants that define or associate with more refined phenotypic subtypes.

Digital pathology has previously been investigated primarily as a stand-in for molecular information when it is unavailable, with genomic sequencing being treated as ground truth^32^. However, clinical next-generation sequencing has important limitations: certain fusions and loss of tumor suppressor function via epigenetic silencing may not be detectable by DNA sequencing, and low-frequency variants can be missed due to measurement depth, panel resolution and variant-calling algorithms^33,34^. Furthermore, many variants in cancer genes have uncertain biologic and clinical significance, with some VUS actually resulting in downstream functional changes^35,36^. This motivates a multimodal approach, incorporating both histopathologic and genomic features for deeper biological profiling^37^.

In this study, we set out to model granular tumor subtypes in a machine-learning-derived embedding space from H&E WSIs and identify genomic variants^38^ and clinically-actionable biomarkers associated with tumor phenotypes. We focus here on a real-world cohort of over 70,000 patients with targeted MSK-IMPACT DNA sequencing ^39^ and matched H&E WSIs. Tumors were annotated with OncoTree^1^, an open-source, community-driven, cancer classification system developed at Memorial Sloan Kettering Cancer Center (MSK) and adopted by the American Association for Cancer Research (AACR) Project Genomics Evidence Neoplasia Information Exchange (GENIE)^40^, to formalize cancer subtyping and associated genomic alterations. We developed a machine learning model, Aeon (adaptive embedding ontology network), to infer OncoTree subtypes from H&E WSIs, finding a highly semantically relevant embedding space through performance testing on internal and external test sets, as well as benchmarking refinement of CUPs against genome-based tools. We also developed models, Paladin (Phenotypic Associative LeArning from Digital pathology Images using Nuanced subtypes) to infer genomic features from H&E WSIs, conditioned on granular subtype and site of disease. We then deployed Paladin to aid with phenotype-based functional annotation of VUS, validating functional significance with outcomes and immunohistochemistry (IHC), and identified histologic phenocopies in two genes, *STK11* ^41,42^ and *FGFR3* ^43,44^. We plan to make this suite of models, collectively called Mosaic, available for public academic use.

## Results

### Real world cohort with matched H&E and clinical genome sequencing

We constructed a pan-cancer real-world cohort of 378,123 H&E WSIs and molecular data (**Fig. 1a**) from 79,149 tumors across 71,142 patients (**Supp. Table 1**) spanning 163 OncoTree-defined cancer subtypes (**Supp. Table 2**)^1^. WSIs from the formalin-fixed, paraffin-embedded parts used for sequencing and OncoTree code assignment (**Fig. 1a**) were tiled at 20x magnification using custom open-source software, Mussel (**Methods**), and pre-processed to exclude low-quality tissue. This yielded 4.74 billion 112 µm x 112 µm tiles for analysis. At the most granular OncoTree code available (with additional stratification by MSI status for stomach, colorectal, and endometrial cancers and hormone/Her2 receptor subtypes for breast cancer), we divided patients into 70% training, 20% test, and 10% validation sets. We trained a model, Aeon, to associate OncoTree codes with H&E WSIs. Within each subtype, we trained Paladin, a set of models to infer genomic targets. Primary and metastatic specimens were tested separately to avoid confounding. Binary genomic targets were considered when at least 50 samples each with and without the alteration were available for each OncoTree code and sample type; continuous-valued targets, such as fraction of genome altered, were included when at least 50 samples were available.

**Figure 1.**
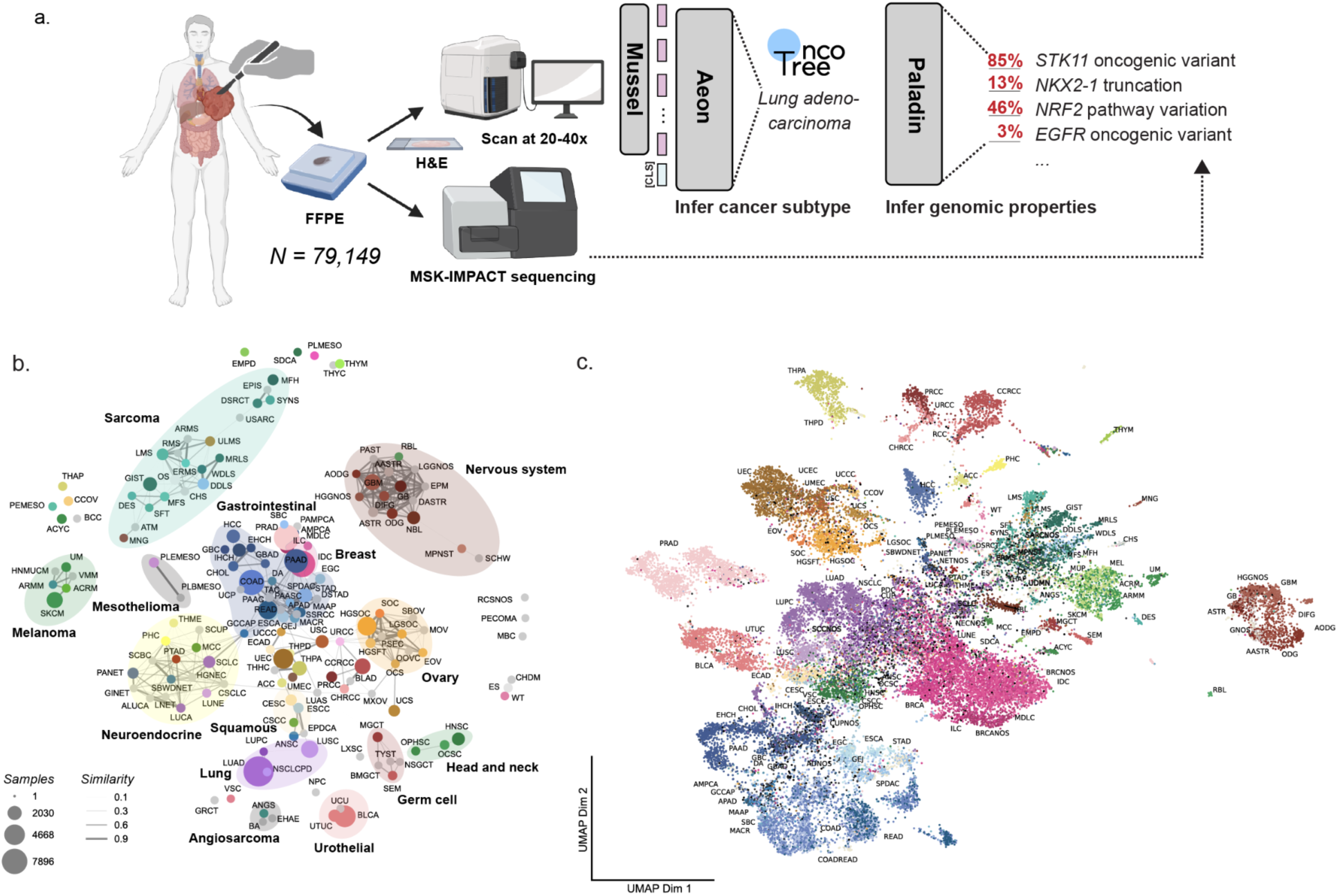
Schematic overview. (a) Formalin-fixed, paraffin-embedded (FFPE) tissue from resected primary or metastatic specimens was used to generate hematoxylin and eosin (H&E)-stained whole-slide images (WSIs) and MSK-IMPACT targeted clinical sequencing results. A statistical learning pipeline preprocesses WSIs using Mussel, infers OncoTree code using Aeon, and infers genomic variants using subtype-specific sub-models of Paladin. (b) Learned graph of OncoTree histologic subtypes, with vertex size determined by log relative abundance and edge weight determined by distance in concept space. (c) UMAP for the test set based on H&E WSIs, colorized by pathologist-labeled OncoTree code. Panel (a) partially created in BioRender by Boehm, K. (2025) [https://BioRender.com/qmhoe64].

**Table 1.**
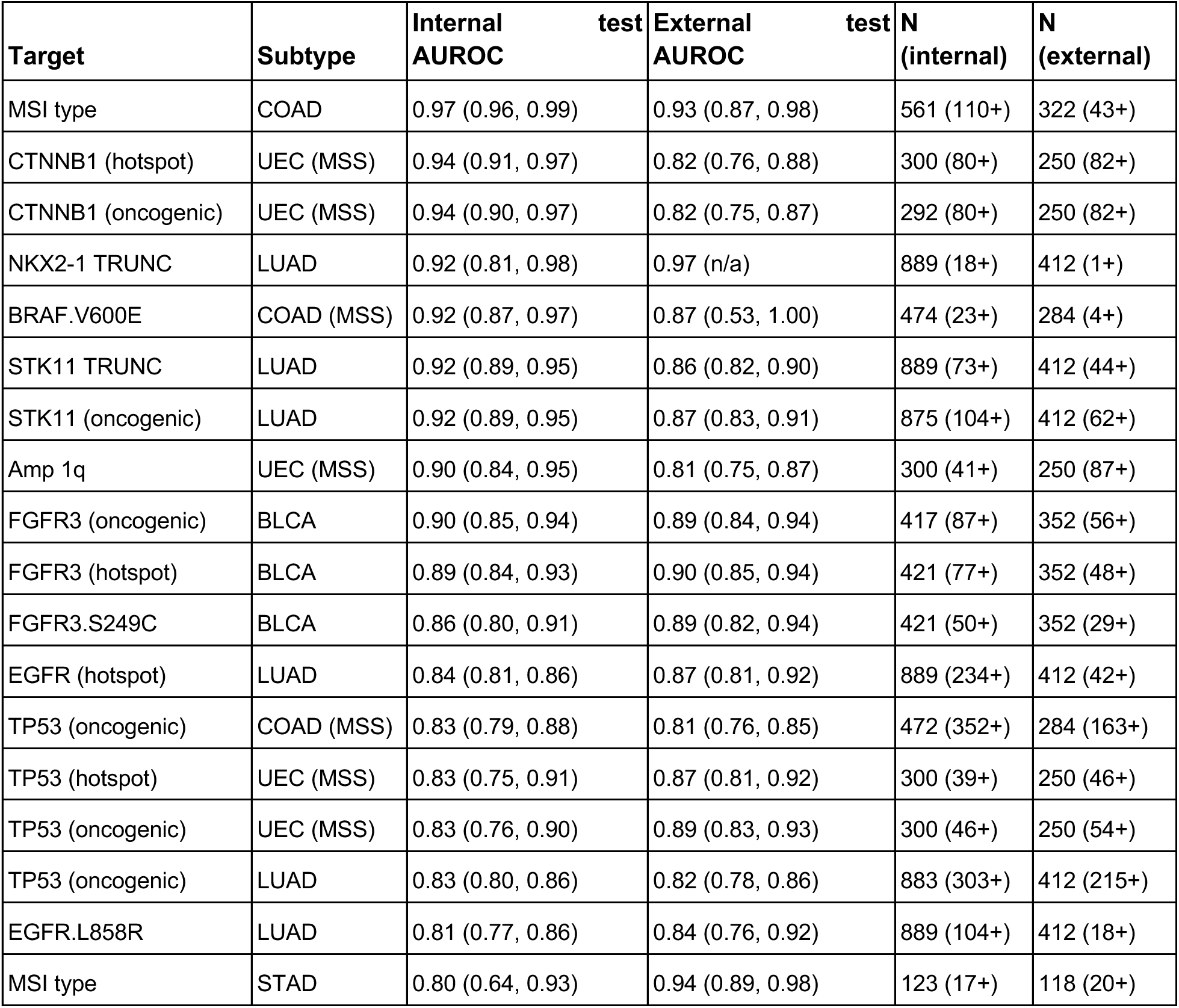
Selected performance metrics. Area under the receiver operating characteristic curve (AUROC) values with 95% confidence intervals by 1000-fold bootstrapping in parentheticals for primary specimens with AUROC ≥ 0.80 on both the internal and external test sets. COAD: colon adenocarcinoma, UEC: uterine endometrioid carcinoma, MSS: microsatellite stable, LUAD: lung adenocarcinoma, BLCA: urothelial carcinoma, STAD: stomach adenocarcinoma.

| Target | Subtype | Internal AUROC<br>test | External AUROC<br>test | N<br>(internal) | N<br>(external) |
| --- | --- | --- | --- | --- | --- |
| MSI type | COAD | 0.97 (0.96, 0.99) | 0.93 (0.87, 0.98) | 561 (110+) | 322 (43+) |
| CTNNB1 (hotspot) | UEC (MSS) | 0.94 (0.91, 0.97) | 0.82 (0.76, 0.88) | 300 (80+) | 250 (82+) |
| CTNNB1 (oncogenic) | UEC (MSS) | 0.94 (0.90, 0.97) | 0.82 (0.75, 0.87) | 292 (80+) | 250 (82+) |
| NKX2-1 TRUNC | LUAD | 0.92 (0.81, 0.98) | 0.97 (n/a) | 889 (18+) | 412 (1+) |
| BRAF.V600E | COAD (MSS) | 0.92 (0.87, 0.97) | 0.87 (0.53, 1.00) | 474 (23+) | 284 (4+) |
| STK11 TRUNC | LUAD | 0.92 (0.89, 0.95) | 0.86 (0.82, 0.90) | 889 (73+) | 412 (44+) |
| STK11 (oncogenic) | LUAD | 0.92 (0.89, 0.95) | 0.87 (0.83, 0.91) | 875 (104+) | 412 (62+) |
| Amp 1q | UEC (MSS) | 0.90 (0.84, 0.95) | 0.81 (0.75, 0.87) | 300 (41+) | 250 (87+) |
| FGFR3 (oncogenic) | BLCA | 0.90 (0.85, 0.94) | 0.89 (0.84, 0.94) | 417 (87+) | 352 (56+) |
| FGFR3 (hotspot) | BLCA | 0.89 (0.84, 0.93) | 0.90 (0.85, 0.94) | 421 (77+) | 352 (48+) |
| FGFR3.S249C | BLCA | 0.86 (0.80, 0.91) | 0.89 (0.82, 0.94) | 421 (50+) | 352 (29+) |
| EGFR (hotspot) | LUAD | 0.84 (0.81, 0.86) | 0.87 (0.81, 0.92) | 889 (234+) | 412 (42+) |
| TP53 (oncogenic) | COAD (MSS) | 0.83 (0.79, 0.88) | 0.81 (0.76, 0.85) | 472 (352+) | 284 (163+) |
| TP53 (hotspot) | UEC (MSS) | 0.83 (0.75, 0.91) | 0.87 (0.81, 0.92) | 300 (39+) | 250 (46+) |
| TP53 (oncogenic) | UEC (MSS) | 0.83 (0.76, 0.90) | 0.89 (0.83, 0.93) | 300 (46+) | 250 (54+) |
| TP53 (oncogenic) | LUAD | 0.83 (0.80, 0.86) | 0.82 (0.78, 0.86) | 883 (303+) | 412 (215+) |
| EGFR.L858R | LUAD | 0.81 (0.77, 0.86) | 0.84 (0.76, 0.92) | 889 (104+) | 412 (18+) |
| MSI type | STAD | 0.80 (0.64, 0.93) | 0.94 (0.89, 0.98) | 123 (17+) | 118 (20+) |

### Aeon identifies 163 granular cancer subtypes alongside major genomic phenotypic determinants

We first investigated the capacity to assign 163 OncoTree granular subtypes using H&E WSIs. We trained the Aeon model using an ontology-smoothed loss function based on knowledge graphs (**Methods**) to learn the weighted graphical relation of OncoTree codes based on biological similarity (**Fig. 1b**). We examined the semantic structure of the embedding (one embedding per part) space by ground-truth, clinically labeled histology (**Fig. 1c**). To probe the determinants of semantic structure, we performed Leiden clustering ^45^ on the whole-part representations (**Fig. 2a**) and observed separation of cancer subtypes (excluding the training set) across 104 Leiden clusters, with adjusted Rand scores by OncoTree code of 0.431 (95% C.I. 0.425-0.439) and 0.320 (95% C.I. 0.306-0.338) for primary and metastatic specimens, respectively (**Fig. 2a**). This supports partitioning of the learned representation space by histologic subtype, especially for primary specimens. To formally probe the separation of OncoTree codes by these clusters, we tested for enrichment of individual OncoTree codes using Fisher’s Exact Test, finding that all OncoTree codes except three (undifferentiated carcinoma of the pancreas, other ovarian cancer, and bladder adenocarcinoma) were enriched in specific clusters at a 1% controlled false discovery rate (FDR; **Supp. Table 3; Extended Data Fig. 1**). We also tested for sub-clustering of microsatellite instability and sample type, identifying that these features were associated with whole-part representations in some cancer subtypes (**Supp. Table 4**).

**Figure 2.**
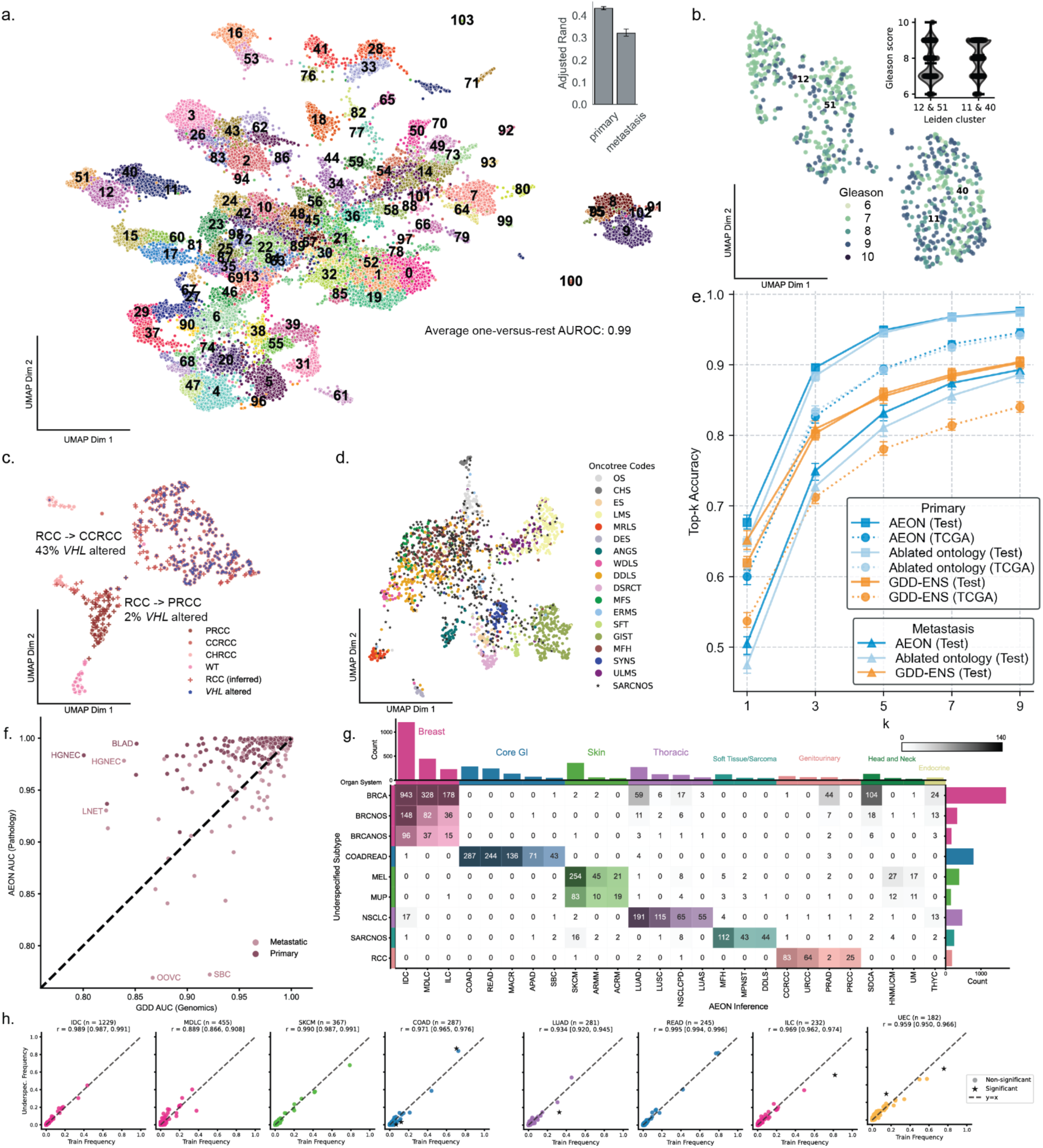
Evaluating histopathologic representations learned by Aeon. (a) UMAP for the test set based on H&E WSIs, colorized by Leiden cluster identity. (b) UMAP for primary prostate adenocarcinoma specimens colorized by pathologist-labeled Gleason score. Enrichment for high Gleason Score was significant across clusters (Mann-Whitney U = 47597.0, *p*-value= 2.8e-11). (c) UMAP for renal malignancies colorized by ground-truth histology (dots) or nearest-neighbor based reassignment (+ marks). (d) UMAP for sarcoma specimens. (e) Top-k accuracy for Aeon and GDD-ENS, with ablation of knowledge graph-based training. (f) One-versus-rest test AUROC values by histology, stratified by site, for AEON vs GDD-ENS, a genome-based classifier. (g) Reclassified underspecified specimens (test set). (h) Genomic concordance analysis between oncogenic variants in training instances and AEON-reclassified underspecified samples, n = number of underspecified instances inferred as each subtype.

We next analyzed how the granularity of Aeon embeddings related to key covariates. For prostate adenocarcinoma primary specimens, we tested the association of Gleason score with Leiden cluster membership, finding that clusters 11 and 40 exhibited higher Gleason score than clusters 12 and 51 (**Fig. 2b**; Mann-Whitney U = 47597.0, *p*-value= 2.8e-11). For kidney cancers, we harnessed the semantic value of the representation space to reclassify underspecified histologies labeled generally as renal cell carcinoma (RCC), which co-segregated with papillary, clear cell, chromophobe, and Wilms tumor specimens, using nearest-neighbor classification. 43% of those reassigned as clear cell exhibited alterations in *VHL*, compared to 2% for those reassigned as papillary (**Fig. 2c**). Similarly, chromosomal arm 3p was deleted in 35% of cases reassigned as clear cell and 11% of cases reassigned as papillary. In sarcomas (**Fig. 2d**) we observed a manifold characterized by poorly differentiated disease in the center, comprising high-grade spindle cell sarcoma and myxofibrosarcoma, branching into increasingly more differentiated disease: chondrosarcoma into osteosarcoma along one axis, uterine leiomyosarcoma into non-uterine leiomyosarcoma along another, and dedifferentiated liposarcoma into well-differentiated liposarcoma along a third axis. Together, these findings exemplify robust representation of granular histologic subtypes from pathology images and further refinement associated with molecular covariates.

We next investigated Aeon performance in a supervised fashion on the held-out test set of 11,693 samples corresponding to the 163 OncoTree codes. Overall AUROC for Aeon was 0.992 on this set with all subtypes AUROC ≥ 0.80, and 161/163 subtypes with AUROC ≥ 0.90 (only “ovarian cancer, other” at 0.82 and “small bowel cancer” at 0.89 had test AUROC < 0.90; **Supp. Table 5**). Despite the complexity of training across 163 detailed subtypes, top-one accuracy for all Aeon outputs reached 62% (top-two: 78%; top-three: 85%; **Fig. 2e**). Ablating the knowledge graph-based training paradigm reduced performance (**Fig. 2e**), e.g., from 0.51 (95% C.I. 0.49-0.51) to 0.47 (95% C.I. 0.46-0.49) top-one accuracy in the test set. Current histology-based subtyping models only used 18 broad cancer labels, which aggregate up to 13 distinct OncoTree codes into a single type (e.g., grouping melanoma, cutaneous squamous cell, basal cell, and merkel cell carcinomas as ‘skin cancer’). Re-mapping these granular OncoTree codes to the corresponding coarse groups resulted in a top-one accuracy of 87%. We also benchmarked Aeon’s performance against a previously developed genomic model, specifically a hyperparameter ensemble model, GDD-ENS^6^, the state of the art for subtype classification from MSK-IMPACT data. We used the same cohort and a slightly adapted GDD-ENS architecture and feature set to re-train GDD-ENS to output the 163 OncoTree code labels learned by Aeon (**Methods**).

The overall AUROC for GDD-ENS was 0.986, with AUROC ≥0.90 for 150/163 subtypes. Notably, 156/163 subtypes had higher AUROC with Aeon compared to GDD-ENS, although the difference was modest with a mean increase per subtype of 0.025. The benefit of the pathology-based approach was slightly reduced when we accounted for sample type: while primary instances had higher AUROC in Aeon for 157/163 subtypes, only 85/163 showed higher performance in the metastatic setting (**Supp. Table 6**), presumably owing to background tissue of primary niches. We compared type-specific AUROC for each subtype with at least five test examples in **Fig. 2f**, highlighting the top two subtypes via AUROC differential per site by either model, which revealed modality-induced performance improvements (e.g., metastatic ovarian cancer, other (OOVC) in genomics, primary bladder adenocarcinoma (BLAD) in pathology). Overall, 43.4% of test samples were correctly inferred by both models, 19.7% were incorrectly inferred by both models, with 18.4% correct in Aeon only and 18.5% by GDD-ENS only, highlighting the complementary nature of H&E WSI and genomic approaches. For example, in the rare subtype of ovarian granulosa cell tumors (GRCT; **Extended Data Fig. 2**), we found that of the 11 pathologist-annotated GRCT specimens in the test cohort, Aeon inferred eight as GRCT, all of which exhibit the characteristic *FOXL2* alteration. Of the three that Aeon did not label as GRCT (labeled as myxoid ovarian cancer (MXOV) for two and uterine sarcoma (USARC) for the other one), two lacked *FOXL2* alterations, and two were also labeled as other subtypes by GDD-ENS (invasive ductal carcinoma (IDC) and leiomyosarcoma).

In general, cancer subtypes with larger training sizes and higher proportions of primary samples outperformed types with smaller training sizes and higher metastatic sample representation (**Extended Data Fig. 3a-b)**. Aeon nevertheless achieved high accuracy for several rarer subtypes, e.g. Schwannoma (SCHW) and Ependymoma (EPM) with training N < 40, as well as majority-metastatic subtypes, e.g., Medullary Thyroid Cancer (THME), Cutaneous Melanoma (SKCM) with >65% metastatic samples. Aeon also achieved high performance across all represented organ systems (**Extended Data Fig. 3c)**, altogether indicating Aeon is able to provide accurate granular histology inference across a wide variety of cancer subtypes.

Aeon misclassifications were informative in the context of the target subtype. The top incorrect target-inference pairs were often within the same organ system, either between highly related or overlapping subtypes, (e.g. true glioblastoma multiforme (GBM) inferred to be generic glioblastoma (GB)) or those known to be histopathologically similar (e.g., true colon adenocarcinoma (COAD) inferred to be mucinous adenocarcinoma of the colon/rectum (MACR) (**Extended Data Fig. 3d)**. This was further confirmed after re-selecting the inferred organ system after averaging the post-softmax logits per system. Overall, 66% of incorrect inferences were classified within the correct organ system by top-one inference, and 68% by logit-mass, with the only substantial cross-system signal related to morphologically similar subtypes in separate organs (e.g. peritoneal and pleural mesothelioma). To counter this, we also estimated inference correctness via Aeon’s biological similarity scores. Mapping these distances to test set target-inference pairs found that mean similarity score across incorrect inferences was 0.77, with 31% of cases with ≥ 0.9 similarity (c.f. random chance mean = 0.60, paired t-test *p* <.0001, Cohen’s d = 0.732; **Extended Data Fig. 4a)**, suggesting the embedding space still conveys relevant information content even with incorrect classifications. Overall, Aeon inferences were highly accurate across diverse and granular cancer type inputs, with incorrect inferences still providing information content relevant to ground truth histology.

To complete interpretation of Aeon performance, we next analyzed 6,018 underspecified cases, where broad histologies were recorded (e.g., invasive breast carcinoma (BRCA)), but detailed subtypes were not (e.g., IDC). We hypothesized Aeon inference would provide further subtype specification when not available or determinable following pathologist review. Of these cases, 77.3% of cases were predicted in the same organ system as the initial underspecified subtype, with the highest recall within the Core GI and CNS systems (94% and 92%, respectively). We visualized the underspecified cases across detailed subtypes with n≥25 inferred samples, grouped by organ system (**Fig. 2g**). Which confirms the high organ system recall, with the highest cross-system overlap between BRCA and Salivary Ductal Carcinoma (SDCA), which are known to be histopathologically similar ^46^. Furthermore, underspecified cases exhibited genomic associations similar to their inferred subtypes: oncogenic alteration frequencies in training samples versus underspecified cases among the eight most prevalent Aeon-inferred subtypes demonstrated strong correlation (**Fig. 2h**). Only 7/2835 tested alterations showed discordance between training and Aeon-inferred underspecified cases, often still exhibiting subtype-specific genomic trends consistent with the inferred type. For example, while the frequency of *CDH1* alterations was significantly lower in Aeon-inferred ILC cases than ground-truth cases, *CDH1* was altered significantly more frequently in Aeon-inferred ILC compared to Aeon-inferred IDC. Aeon-inferred groups also recapitulated other genomic trends expected from prior studies (i.e. *TP53*, *GATA3* in IDC, and *PIK3CA*, 16q deletions in ILC ^47^**; Extended Data Fig. 4b**).

Together, these findings support that a semantically rich representation of tumor histopathology was learned by Aeon, shown via unsupervised analysis of the learned space via Leiden clustering and via supervised classification. The model identifies subtype as well as a genomic classifier, including with rare subtypes, and refines coarse diagnoses as corroborated by orthogonal molecular evidence.

### Inference and re-assessment of Cancers of Unknown Primary

We next assessed Aeon inferences on 1,171 samples of CUP or related, unknown subtypes (i.e., CUP not otherwise specified (CUPNOS), adenocarcinoma NOS (ADNOS), squamous cell carcinoma NOS (SCCNOS) and undifferentiated malignant neoplasm (UDMN). Samples were reassigned to 117 distinct subtypes, 10 of which had at least 25 cases reassigned to them (**Fig. 3a**), most commonly gallbladder cancer (GBC, n=105), lung adenocarcinoma (LUAD, n=73) and cholangiocarcinoma (CHOL, n=60). While CUP cases by definition lack ground-truth sites of origin, certain CUP subtypes have histological specifications that describe the expected true cancer subtype, e.g., ADNOS or SCCNOS. For the 455 cases of either ADNOS or SCCNOS within the CUP set, 237/358 ADNOS samples were inferred adenocarcinoma subtypes (e.g., Pancreatic Adenocarcinoma, LUAD), 79/97 SCCNOS samples were inferred as Squamous subtypes (e.g. cervical or lung squamous cell carcinoma (LUSC)), and only 12/455 were inferred across broad types (e.g. ADNOS inferred LUSC), with the remaining cases assigned subtypes without broad tissue determinants (i.e. GBC; **Fig. 3a**).

**Figure 3.**
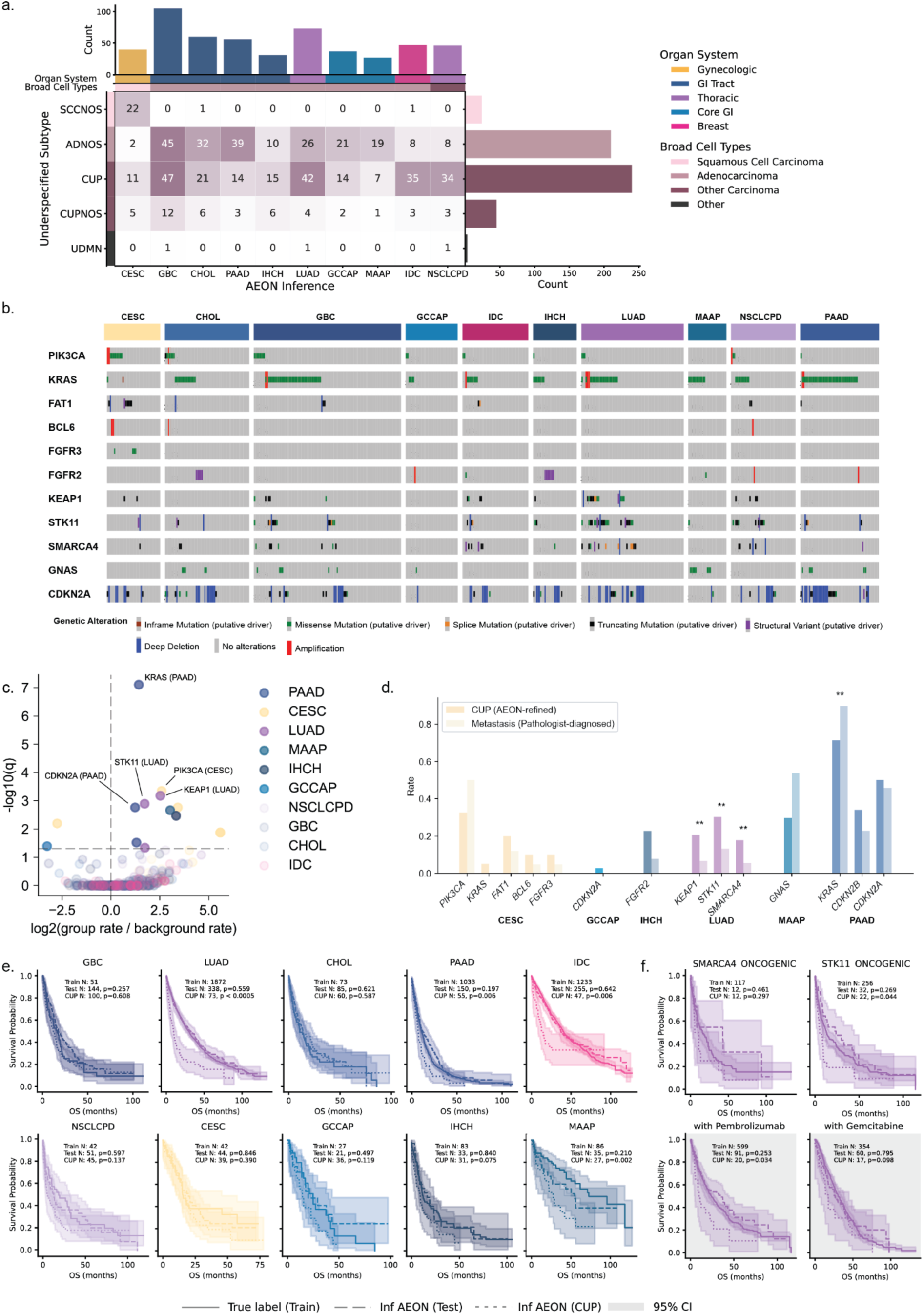
Subtyping cancers of unknown primary. (a) Fraction of AEON-reassigned subtypes by CUP subgroup. (b) OncoPrint for AEON-reassigned groups. (c) Enrichment of genomic variants among AEON-reassigned groups (*p*-value by Fisher’s exact test (one versus rest), false discovery rate controlled by Benjamini-Hochberg method). (d) For variants enriched in at least one group, comparison of abundance of those variants to metastatic specimens of the same histology (*p*-value by Fisher’s exact test, false discovery rate controlled by Benjamini-Hochberg method; transparent points failed to reach significance). (e) Overall survival of AEON-reassigned cases compared to metastatic specimens labeled by pathologists. (f) Granular analysis of survival differences in discrepant cases.

Aeon inferences captured expected trends in the common genomic features of the top-inferred subtypes, such as *FGFR2* fusions in cholangiocarcinomas and *KEAP1*/*STK11* alterations in LUAD (**Fig. 3b**). Concretely, comparing the frequency of OncoKB-annotated variants across Aeon-defined subtypes for CUPs using Fisher’s exact test (**Fig. 3c**), we found that 13 genes were enriched/depleted with 5% FDR. We compared the alteration rates in these genes in the Aeon-defined subtypes against those in pathologist-annotated metastatic specimens (where genomic alteration rates differed among baseline groups; see **Supp. Table 7**), finding that 9/13 did not differ significantly at 5% FDR (**Fig. 3d**). Alterations indicating poor prognosis in LUAD (*KEAP1, STK11, SMARCA4*), with *SMARCA4* previously associated with dedifferentiation^48^, were more common in specimens labeled as CUP by pathologists than those labeled as LUAD. In pancreatic adenocarcinoma (PAAD), *KRAS* alterations were more common in pathologist-labeled PAAD than CUPs inferred to be PAAD by Aeon.

We next determined the model’s prognostic value by comparing overall survival (OS) across Aeon-assigned subtypes to OS of true metastatic samples of each subtype (which exhibited different OS at baseline with log-rank p=2.7e-136). OS was not significantly different between train samples and Aeon-inferred, metastatic test samples for any of the 10 cancer types highlighted in **Fig. 3a**. Only LUAD showed a significantly different OS in the Aeon-inferred CUP specimens when compared to training (**Fig. 3e**; Bonferroni-corrected p < 0.001). Controlling for the presence of known LUAD-related oncogenic alterations and common LUAD treatment regimens reduced this differential signal in the affected group (**Fig. 3f**, **Extended Data Fig 5**). Thus, the Aeon embedding space has potential to enable refinement of CUPs into granular subtypes.

### Genomic features exhibiting tumor subtype-specific characteristic phenotypes on H&E

We next investigated the capacity to infer genomic information based on corresponding H&E WSIs, conditioned on the granular OncoTree tumor subtypes. We began in an unsupervised setting by assessing which oncogenic genomic events were associated with Aeon Leiden cluster membership. We tested this within a specific histology, limited to only primary or metastatic samples (**Fig. 4a**) and microsatellite stable disease. Seventy genomic features were enriched by Leiden cluster membership with 1% controlled FDR (**Fig. 4a, Supp. Table 8**). Top examples by odds ratio were *TERT* mutational status in primary urothelial carcinoma of the urinary bladder (BLCA; **Extended Data Fig. 6a**), *BRAF* mutational status in primary papillary thyroid cancer (**Extended Data Fig. 6b**), *TP53* and *EGFR* mutational status in primary lung adenocarcinoma (**Extended Data Fig. 6c-d**), *TERT* mutational status in primary glioblastoma (**Extended Data Fig. 6e**), *TP53* mutational status in primary invasive ductal carcinoma of the breast (**Extended Data Fig. 6f**), *EGFR* mutational status in metastatic lung adenocarcinoma (**Extended Data Fig. 6g**), and *KRAS* mutational status in primary lung adenocarcinoma (**Extended Data Fig. 6h**). To assess effect size, we calculated Cramer’s V value (**Extended Data Fig. 7**) and found that *TP53* (established determinant of tumor grade and differentiation ^49^), was a major driver of phenotypic variation across multiple histologies and sample types, as were *RB1* (a known driver of neuroendocrine differentiation ^50^), *CDKN2A*, *KRAS*, *APC*, *TERT*, and *EGFR* (**Extended Data Fig. 7**).

**Figure 4.**
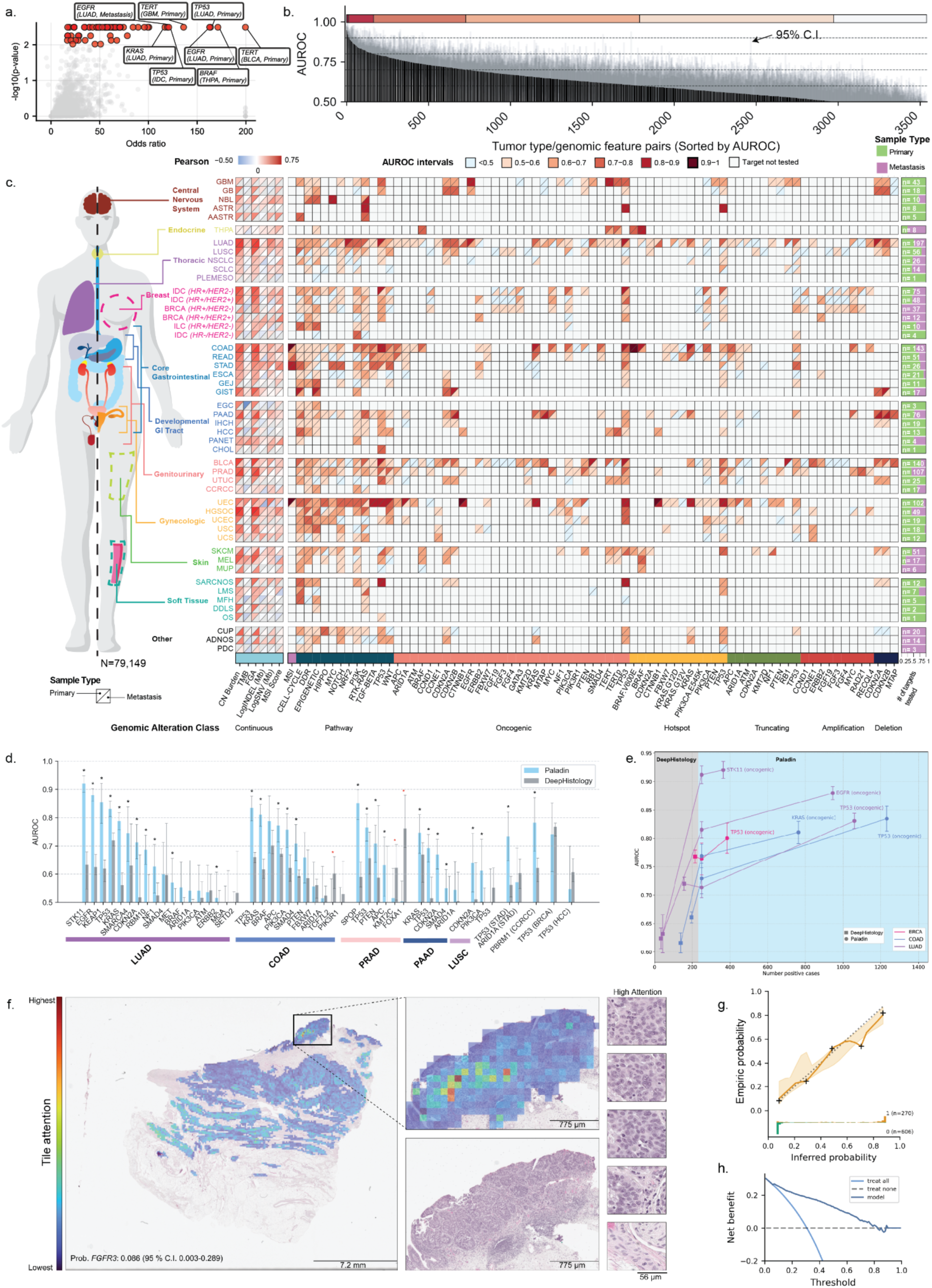
Cancer subtype-specific inference of genomic features by Paladin. (a) Volcano plot of Fisher’s exact test odds ratios and 1% false discovery rate-controlled *p* values for associations of oncogenic mutations in a given gene with Leiden cluster in Aeon representation space, conditioned on specific OncoTree code and sample type. (b) Test area under the receiver operating characteristic curve (AUROC) values with 95% confidence interval (grey). (c) Test AUROC and Pearson correlation values for tested subtype-feature pairs, stratified by primary or metastatic site. (d) Task-specific benchmarking of Paladin against DeepHistology. (e) Data ablation comparison against DeepHistology. (f) Saliency map for inference of *FGFR3* oncogenic variants in urothelial carcinoma specimen with highest-attention tiles shown at higher power. (g) Calibration curve for *EGFR* oncogenic variant inference in lung adenocarcinoma across five quantiles with 95% confidence interval shown by Lowess smoothing using 1000-fold bootstrapping. (h) Net benefit curve for *EGFR* oncogenic variant inference in primary lung adenocarcinoma.

We next trained a series of supervised models, Paladin (**Methods**), to infer actionable biomarkers from H&E. Of 3,541 unique histology-target pairs across 52 granular histologies (**Supp. Table 9,10**), performance of 165 (4.7%) had an AUROC of 0.80 or greater (**Fig. 4b**, **Fig. 4c**). The genomic features most strongly associated with H&E (internal and external test AUROC ≥ 0.80) included expected associations such as MSI in COAD and stomach adenocarcinoma, oncogenic *CTNNB1* alterations in MSS uterine endometrioid carcinoma, *BRAF* V600E alterations in MSS COAD, and *EGFR* oncogenic alterations in LUAD (**Table 1**). On 48 overlapping histology-target pairs Paladin achieved significantly better performance compared to DeepHistology ^29^ in 28 cases (58%), comparable performance in 17 cases (35%), and worse performance in three cases (6%) (**Fig. 4d**). Examining variants previously reported to be associated with phenotype on H&E WSIs ^30^ in coarse cancer subtypes, we compared against subtype-specific performance, finding that *MEN1* and *CDH1* mutational status in aggregated pancreatic and breast cancers were simply associated with the neuroendocrine and lobular subtypes, respectively, rather than characteristic phenotypes within those subtypes (**Extended Data Fig. 8**). For six high-performing models, we ablated training dataset size to 250 positive cases (**Fig. 4e**), showing asymptotically increasing performance with size. Saliency maps were generated to check for confounding (e.g., *FGFR3* in BLCA in **Fig. 4f**), showing tumor cells in the highest-attention tiles. Calibration curves (e.g., *EGFR* in LUAD in **Fig. 4g**) and net benefit curves (**Fig. 4h**)^51^ were generated for each histology-target pair. Analyzing phenotypic associations with genomic alterations at a granular level thus identified new and strengthened genomic associations, while overcoming confounding covariates in previously reported associations analyzed at the level of coarse subtypes.

### Orthogonal Measurements Suggest Informative Disagreement with Sequencing

Next, we set out to systematically clarify the functional effects of VUS. For Paladin models with test AUROC ≥ 0.75, we compared the inferred probability of alteration for cases with missense VUS against wildtype cases using one-sided Mann-Whitney U tests with 5% FDR. Considering missense mutations generating the same amino acid substitution occurring three or more times, we found 22 with greater-than-background inferred probability of oncogenic alteration (**Fig. 5a**). Of these, VUS with Cohen’s d ≥ 2.0 included R234W, L281M, G332C, S224F, and R362Q in *KEAP1* in LUAD along with G332C and F104C in *SPOP* in prostate adenocarcinoma (**Fig. 5b**). These results suggest functional significance of these missense mutations and could help guide annotation databases such as OncoKB based on phenotypic changes observed on routine H&E WSIs. For stage 4 LUAD tumors, we stratified *STK11* VUS into functionally significant (VUS (sig.)) and insignificant (VUS (insig.)) groups with 5% FDR. For these groups, we compared overall survival (OS; **Fig. 5c**) to those with oncogenic or likely oncogenic variants by OncoKB and those with wildtype (WT) tumors. We observed that VUS (sig.) cases exhibited shorter OS compared to WT cases (log-rank p=1.6e-5), and not significantly different from that of cases with oncogenic variants (p=0.08), while VUS (insig.) did not exhibit OS significantly different from wild-type cases with 1% FDR (p=0.020). These disparate outcomes further support H&E phenotype-based functional annotation of VUS using Paladin.

**Figure 5.**
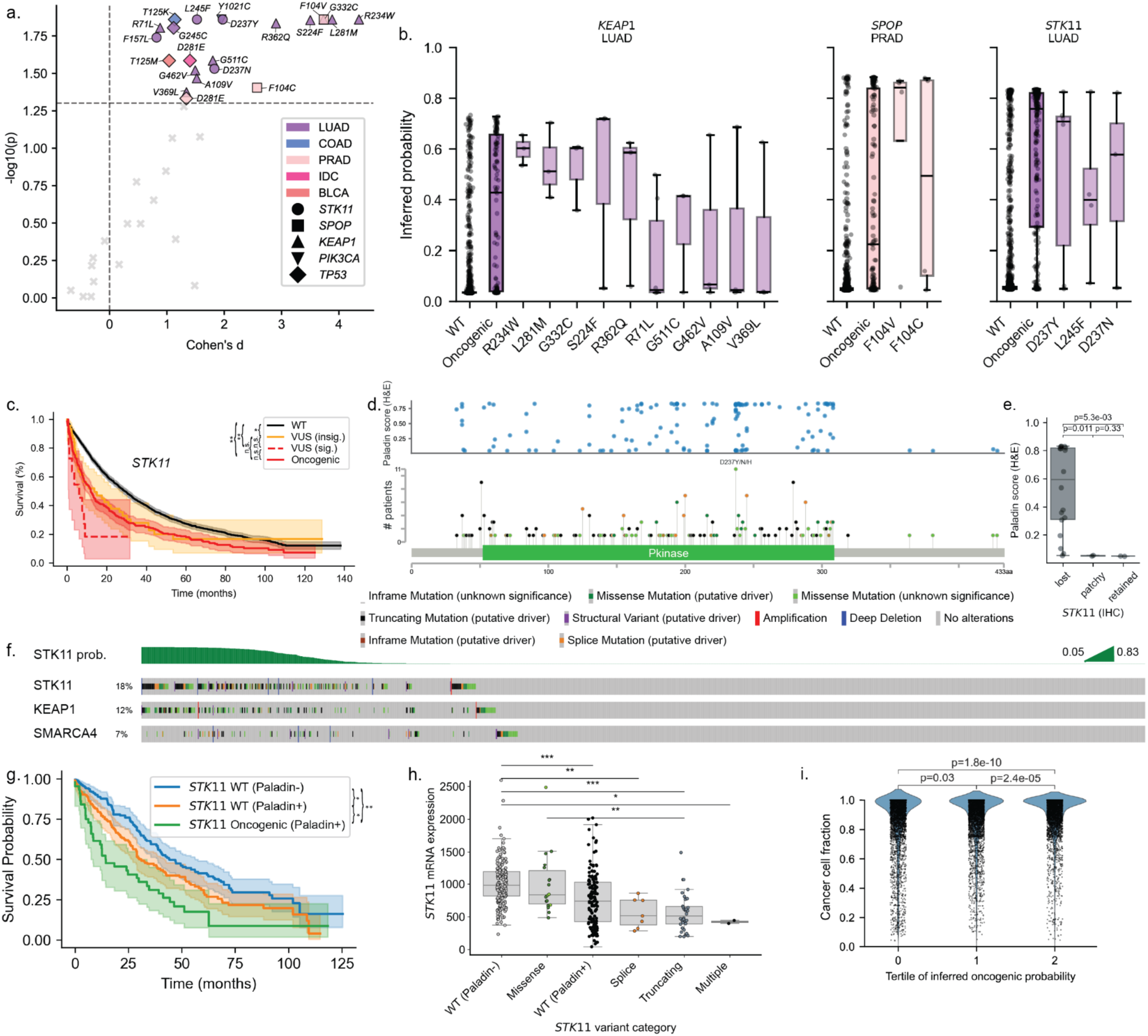
Analysis of variants of uncertain significance, *STK11* phenocopiers, and effect of cancer cell fraction on model confidence. (a) Missense mutations by individual OncoTree code, gene, and sample type (p values by Mann-Whitney U test of Paladin probabilities for cases with missense VUS against the distribution of Paladin probabilities for wild-type cases. (b) Box plots of inferred Paladin probabilities for cases shown in (a) for genes in specific OncoTree codes with at least two VUS reaching significance, regardless of effect size. (c) Overall survival with shaded 95% confidence interval by *STK11* mutational status for stage 4 lung adenocarcinoma, with VUS SNVs with corrected p < 0.05 denoted as significant (sig.) and all others denoted as insignificant (insig.), with survival comparison p values by the log-rank test. (d) Paladin probability of *STK11* oncogenic variant disaggregated by IMPACT-detected variant subtype and locus. (e) Immunohistochemical (IHC) results by Paladin-inferred probability of *STK11* alteration for VUS (n=22). (f) OncoPrint for *STK11* in lung adenocarcinoma. (g) Overall survival with shaded 95% confidence interval by inferred *STK11* mutational status for stage IV cases in the test and validation sets. (h) *STK11* transcription versus inferred probability of oncogenic alteration in the external test set, with p values by the log-rank test. WES: whole-exome sequencing. (i) Cancer cell fraction binned by tertile of inferred probability of oncogenic variant for cases excluding training set with MSK-IMPACT-detected oncogenic variant for models with test AUROC ≥ 0.70. Boxes in (b), (e), and (h) denote interquartile range (IQR), line within denotes median, whiskers denote 1.5 x IQR, and other points are outliers. * denotes p < 0.05, ** denotes p < 0.01.

Lastly, we investigated the cases in which Paladin and MSK-IMPACT disagreed, focusing on *FGFR3* alterations in BLCA and *STK11* alterations in LUAD. We hypothesized that such cases harbored occult molecular alterations that phenocopied the expected impact of genomic alterations. Ordering cases in the test and validation sets by their inferred probability of oncogenic *FGFR3* alteration (**Extended Data Fig. 9a**), most high-probability cases exhibited oncogenic alterations in *FGFR3* or *HRAS.* Alterations annotated as functionally significant by OncoKB were assigned higher probability of significant *FGFR3* alteration than those with VUS (**Extended Data Fig. 9b**). *FGFR3*-altered urothelial cancers had the previously reported ^44^ characteristic phenotype of raisinoid nuclei with papillary morphology (**Extended Data Fig. 9c**) compared to *FGFR3-*WT cases (**Extended Data Fig. 9d**). For WT cases in the external test set, *FGFR3* transcripts were more abundant for cases with higher inferred probability of an *FGFR3* oncogenic variant (Mann-Whitney *q*-value < 0.0001; **Extended Data Fig. 9e**), suggesting phenocopying alterations via epigenetic mechanisms, occult *FGFR3* alterations, or aberrations to other elements of the pathway: for example, we commonly observed *HRAS* alterations for *FGFR3* WT cases with high Paladin probability.

We conducted a similar analysis for *STK11* variants in lung adenocarcinoma, observing that variants of uncertain significance in *STK11* spanned a range of inferred probability scores, suggesting functional significance for some (**Fig. 5d**). We performed IHC assessment of *STK11* function for cases with VUS and found that Paladin probabilities were higher for cases with loss of *STK11* on IHC than those with retained or patchy *STK11* (**Fig. 5d**, Mann-Whitney U p=5.3e-3 and p=0.011, respectively). We also observed *STK11* WT cases with high inferred probability of oncogenic *STK11* alterations (**Fig 5f**), noting expected enrichment of *SMARCA4* or *KEAP1* alterations ^41^. This prompted us to compare prognosis for concordant and discordant cases (**Fig. 5g**): we found that stage 4 cases with *STK11* variants inferred by both expectedly exhibited the shortest OS, cases with *STK11* variants inferred by neither had the longest OS, and WT cases with high Paladin *STK11* scores represented an intermediate prognostic group (log-rank *p*-value = 0.01). For WT cases in the TCGA test set, transcript abundance was lower for cases with high Paladin probabilities (≥0.5) than those with low probabilities (**Fig. 5h**; Mann-Whitney U *q*-value < 0.0001), supporting abrogation of *STK11* function in this intermediate-prognosis group. Taken together, these results support the utility of H&E tumor phenotyping to assess variant significance and identify prognostically significant *STK11* phenocopying groups in LUAD. Finally, we quantified the effect of cancer cell fraction (CCF) on model confidence for Paladin models with test AUROC ≥ 0.70 for cases with oncogenic variants. Confidence scores were slightly higher in cases with higher tumor purity (Spearman’s r=0.08; p=2.5e-13) and CCF (Spearman’s r=0.05; p=6.3e-10), but models performed almost as well in cases with lower CCF (**Fig. 5i**).

## Discussion

In this work, we developed Mosaic, a suite of machine learning models using biological knowledge to learn rich phenotypic representations of tumors from H&E WSIs and analyze drivers of phenotypic variation. Our findings quantitatively establish that embedding spaces derived from H&E WSIs robustly represent granular tumor subtypes reflective of clinical practice, providing a foundation on which to improve diagnosis and disentangle phenotypic associations of biomarkers using routine histopathological images, with immediate clinical relevance. Compared to previous work^10,11^, Mosaic supports an order of magnitude more granular cancer subtypes, advancing from 18 coarsely aggregated cancer types to 163 specific histologies. These models can i) guide difficult diagnoses, such as rare primary histologies or CUPs; ii) identify actionable genomic alterations via H&E phenotypic correlates; iii) improve the annotation of VUS; and iv) identify tumors which lack given genomic alterations but mimic their phenotypic consequences. Modeling granular subtypes not only facilitates state-of-the-art performance on supervised tasks, but also enables unsupervised discovery of phenotypic associations.

We extended digital pathology-based tumor subtyping to the granularity of OncoTree^1^ for application to rare primary histologies and CUPs. Favorable performance compared to state-of-the-art genomic subtyping^6^ establishes digital pathology as an aid during challenging diagnostic processes, even where clinical sequencing is feasible. H&E-based subtyping offers dual advantages of cost savings and shorter turnaround time, with results that can later be supported by clinical sequencing, enabling clinical teams to begin specific therapy or trial enrollment sooner. We thus suggest that digital pathology be incorporated into diagnostic proposal systems to triage specialized diagnostic tests, reducing laboratory assay burden and preserving often-scarce diagnostic tissue. This work also establishes incorporating biological knowledge into black-box digital pathology models, guiding the proposal of biologically-related subtypes. How integrating genomic and histologic features improves diagnostic accuracy in clinical settings where both data modalities are available remains an open question.

Compared to previous work^11,29,52,53^ on the inference of genomic features, the most critical advance is the analysis at the level of granular cancer subtypes. This approach identified that some previously reported^30^ phenotype-genotype associations were merely due to associations with one specific histology over another, such as *CDH1* with lobular histology in invasive breast cancer. Our findings disentangle histologic subtyping from genomic biomarker inference, though the boundary between histopathology-based and molecular-based subtyping is blurred in cases where a genomic feature causes a characteristic tumor appearance. Our approach also demonstrated the capacity to discover new associations beyond supervised tasks. Other advances include making use of a larger training dataset and modern machine learning methods, which significantly improved inference of biomarkers with established phenotypic correlates. These implications can extend more broadly to interpret variants from more comprehensive sequencing, study other classes of somatic genomic alteration such as aneuploidy, genomic instability, and large-scale rearrangements, and study non-genomic properties that impact tumor morphology and cellular architecture, such as epigenetics and the tumor microenvironment.

By integrating orthogonal data modalities, we also established the value of digital pathology-based phenotyping to augment biologic profiling beyond that attained by disaggregated sequencing alone. Considering Mosaic’s role in variant annotation, certain missense VUS (many with conflicting *in vitro* evidence^54–56)^ were consistently inferred to have the same phenotype as those with functionally significant alterations, with prognostic significance and proteomic validation for *STK11* in lung adenocarcinoma. The approach also demonstrated value in identification of histologic phenocopies for *STK11* based on transcriptomic profiling, refining the prognostic beyond that of sequencing alone. Taken together, these results advance digital pathology as an auxiliary approach for functional annotation of VUS in databases such as OncoKB^35^ and identification of prognostically significant phenocopies. We suggest that future work should seek to interpret the histomorphology phenotype associated with loss of *STK11* expression and biological mechanisms underpinning phenocopying states.

Despite a large sample size, the analysis presented here is likely still underpowered to identify associations for rare variants in uncommon cancer types. To infer granular histology, we relied on clinically assigned OncoTree codes at the time of sequencing, which may not have always been maximally specific: to mitigate this, we removed non-specific codes from the training set. Furthermore, we included all H&E WSIs associated with each part used for sequencing and treated each part as a single bag of tiles derived from all associated slides given weakly supervised learning’s robustness to inclusion of uninformative tiles with sufficiently large training datasets^22,57^, but for smaller data subsets for rarer histologies, additional supervision may improve performance. For variant annotation, functionally significant VUS were identified, but further study including pan-cancer variant analysis is warranted. Finally, we trained each model using hyperparameters selected empirically on a paradigmatic task for fair comparison and given computational resource constraints, but performance may be improved by task-specific hyperparameter optimization. Multi-institutional training approaches such as federated learning may further reduce generalization error.

In summary, we have established a suite of machine learning models using H&E WSIs to approach granular cancer subtyping, with measurable implications for interpreting genomic and clinical covariates. We significantly improve on the prior state of the art for genomic feature inference and tumor subtyping and will make the resultant models available for public academic use. We propose our models as a quantitative foundation upon which to study and discover genomic and non-genomic properties of granular tumor morphology.

## Code availability

Mussel is available at https://github.com/pathology-data-mining/Mussel. Paladin and Aeon are undergoing additional software engineering and documentation prior to public release.

## Acknowledgements

This work is supported by the Marie-Josée and Henry R. Kravis Center for Molecular Oncology, The Warren Alpert Center for Digital and Computational Pathology, The Halvorsen Center for Computational Oncology, The Fund for Innovation in Cancer Informatics (Spring 2024, Major Grant), Memorial Hospital Translational Research, Ovarian Cancer Research Alliance (OCRA) Collaborative Research Development Grant [648007], and the National Cancer Institute (NCI) Cancer Center through a Core Grant [P30-CA008748] and P01CA275746. We gratefully acknowledge the resources and services of the High-Performance Computing Group and the Cancer Data Science Initiative at Memorial Sloan Kettering Cancer Center.

---

**Extended Data Fig. 1.**
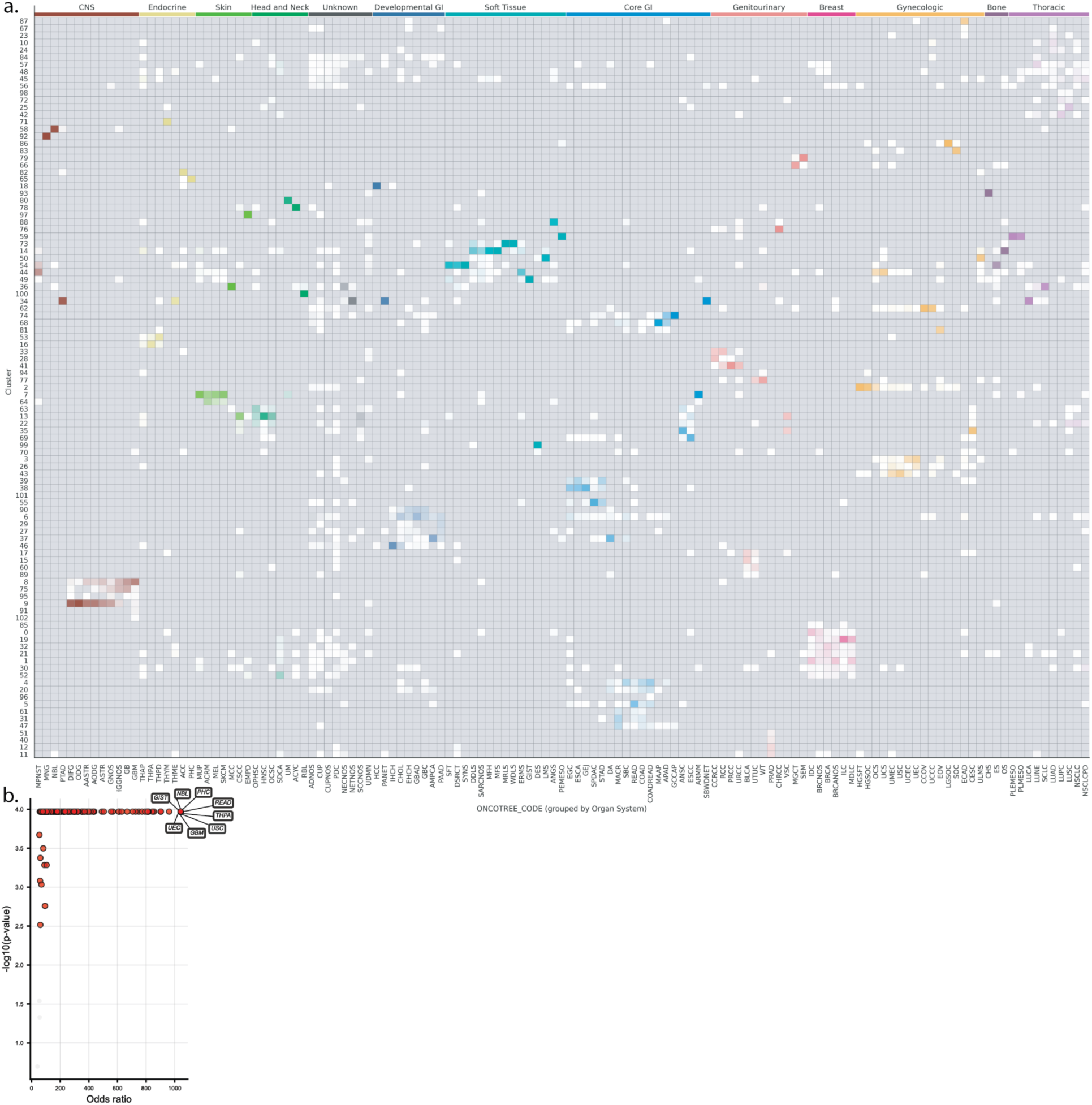
Association of OncoTree codes and oncogenic variants with Leiden clusters. (a) Column-normalized heatmap depicting relative membership of each OncoTree code in each Leiden cluster. (b) Volcano plot showing one-versus-rest enrichment by Leiden cluster as assessed by Fisher’s exact test (controlled for 1% false discovery rate using Benjamini-Hochberg method) for all OncoTree codes tested except three (undifferentiated carcinoma of the pancreas, other ovarian cancer, and bladder adenocarcinoma).

**Extended Data Fig. 2.**
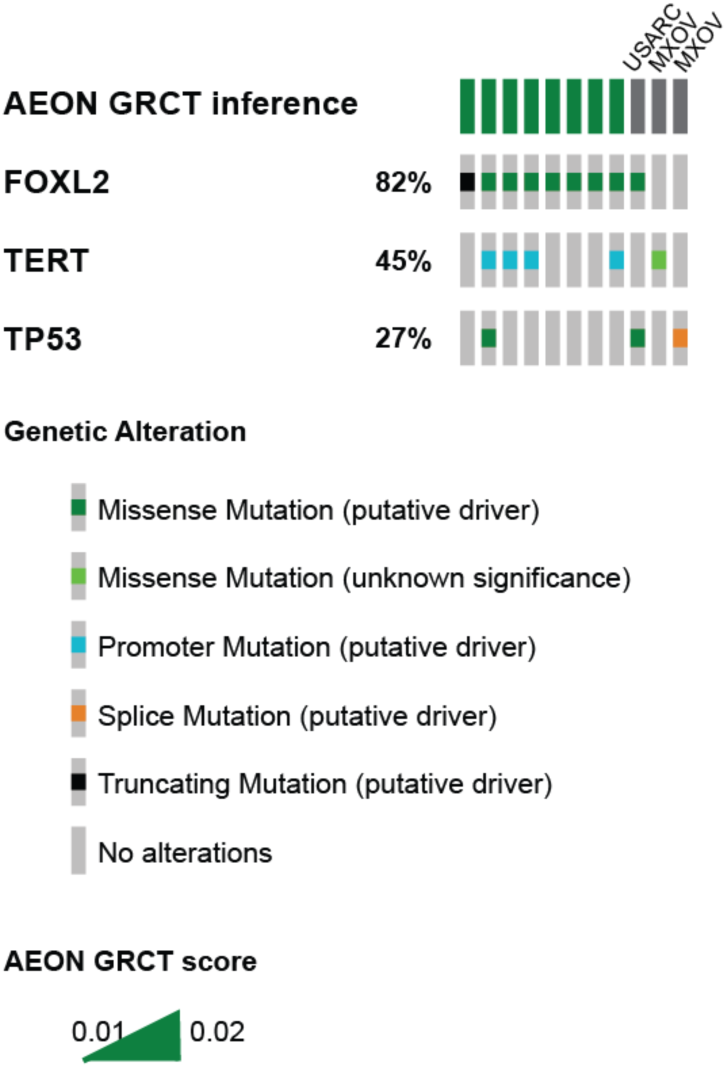
Mutational profile of ovarian granulosa cell tumors in the test set and Aeon-inferred subtype. MXOV: myxoid ovarian cancer, USARC: uterine sarcoma.

**Extended Data Fig. 3.**
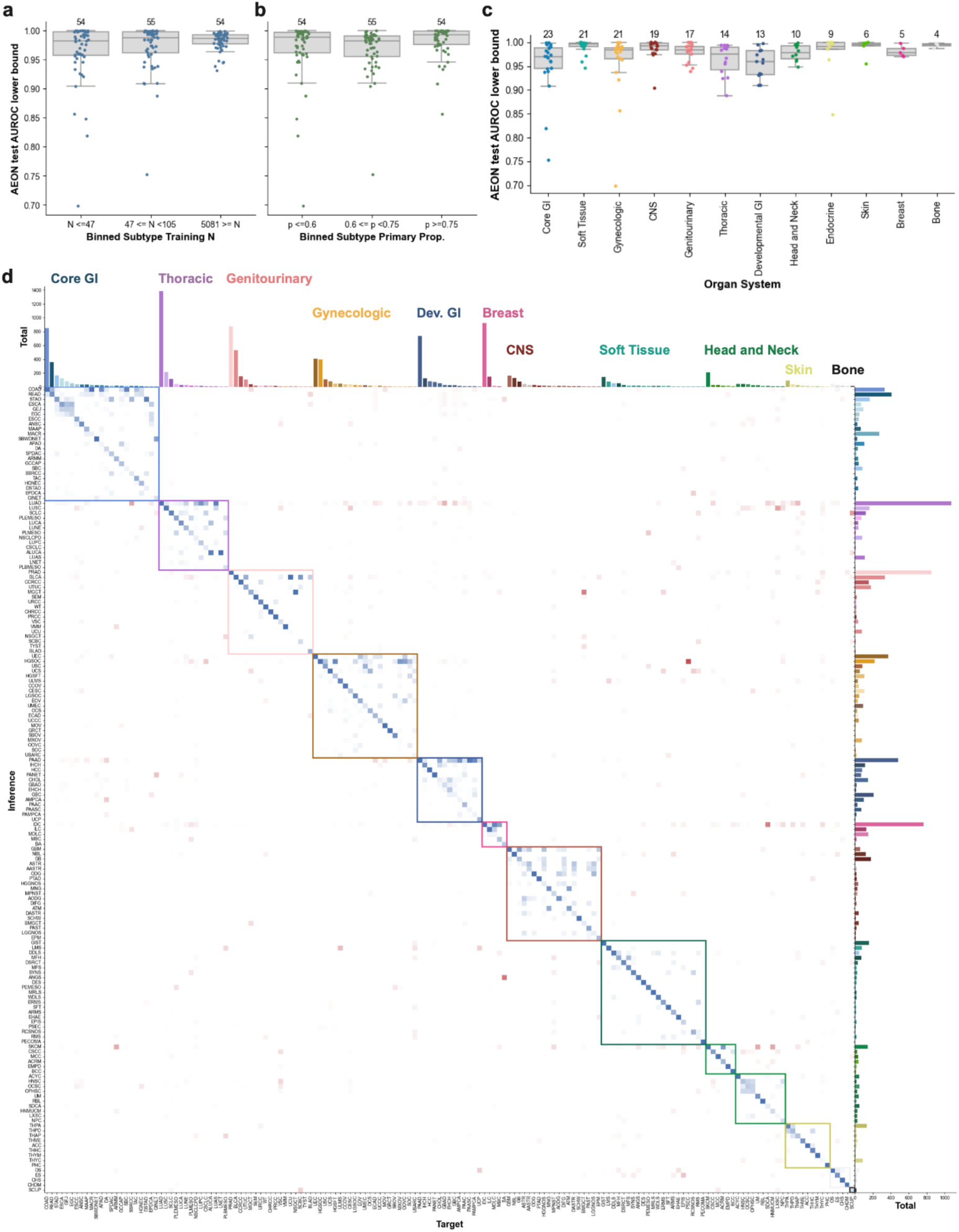
Boxplots showing density and statistics for the lower bound AEON AUROC per subtype after 1000-fold bootstrapping, binned by their training sample size (a) or proportion of primary samples (b) and organ system (c). Numbers above each bar indicate the number of cancer types included in the bin. (d) Column-normalized confusion matrix of AEON inferences grouped by organ system, in order of target organ system and cancer subtype size. Blue indicates inferences within the correct organ system, red indicates cross-system signal, where vmax = 1). Boxes denote interquartile range (IQR), line within denotes median, whiskers denote 1.5 x IQR, and other points are outliers.

**Extended Data Fig. 4:**
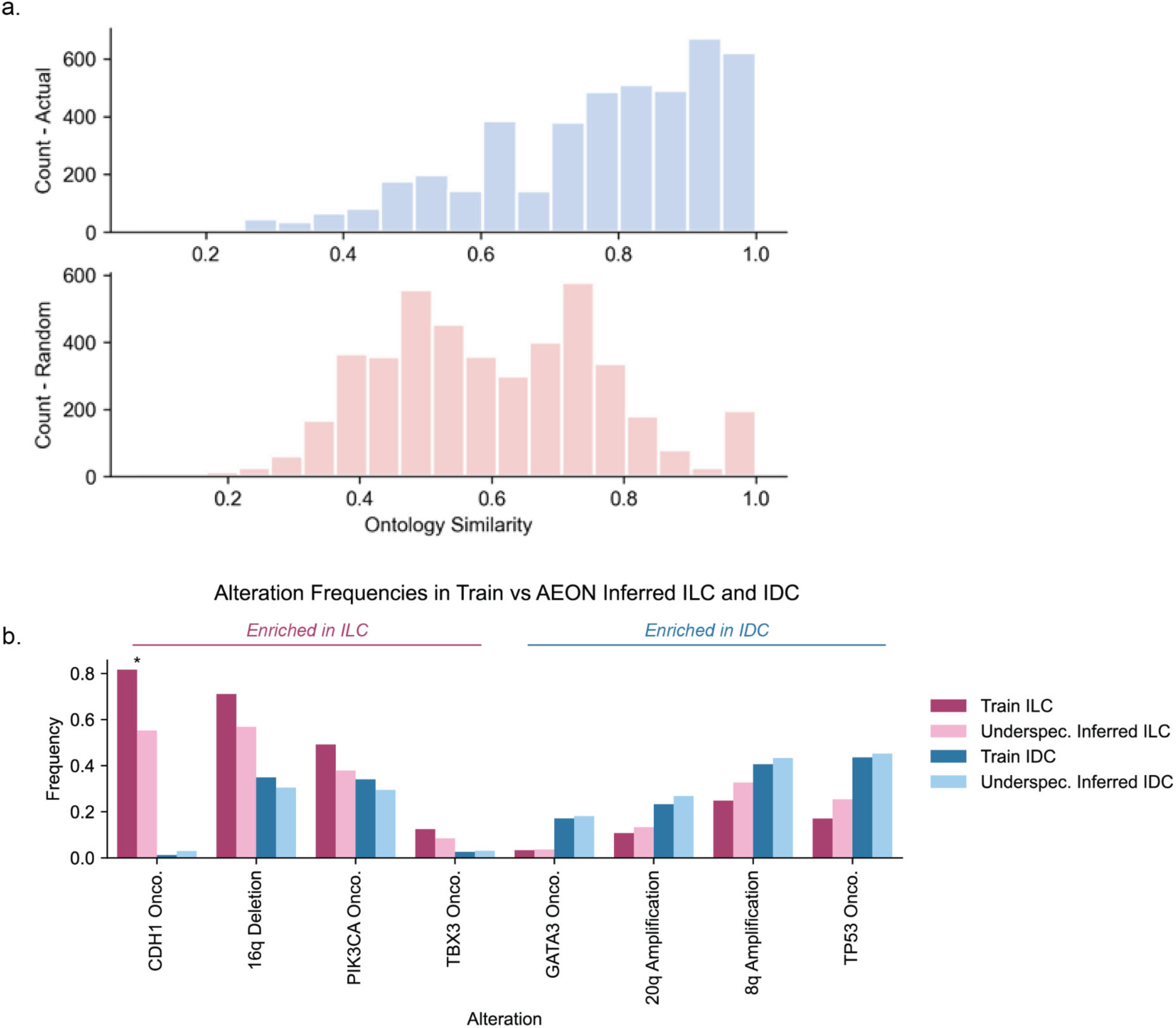
(a) Histogram of ontology similarity values for incorrect AEON test inferences (top) and target-inference pairs generated from the ground truth targets of the incorrect AEON test samples, and an inference label randomly sampled from the distribution of true target types in the full AEON test set, with replacement (bottom). (b) Frequencies of alterations with known, differential genomic trends between IDC and ILC. After comparing frequencies of each alteration between the ground truth, training set cases to AEON-inferred cases within the underspecified set, only CDH1 oncogenic alterations showed a significant underenrichment within the underspecified cases inferred as ILC (*p* = .001).

**Extended Data Fig. 5.**
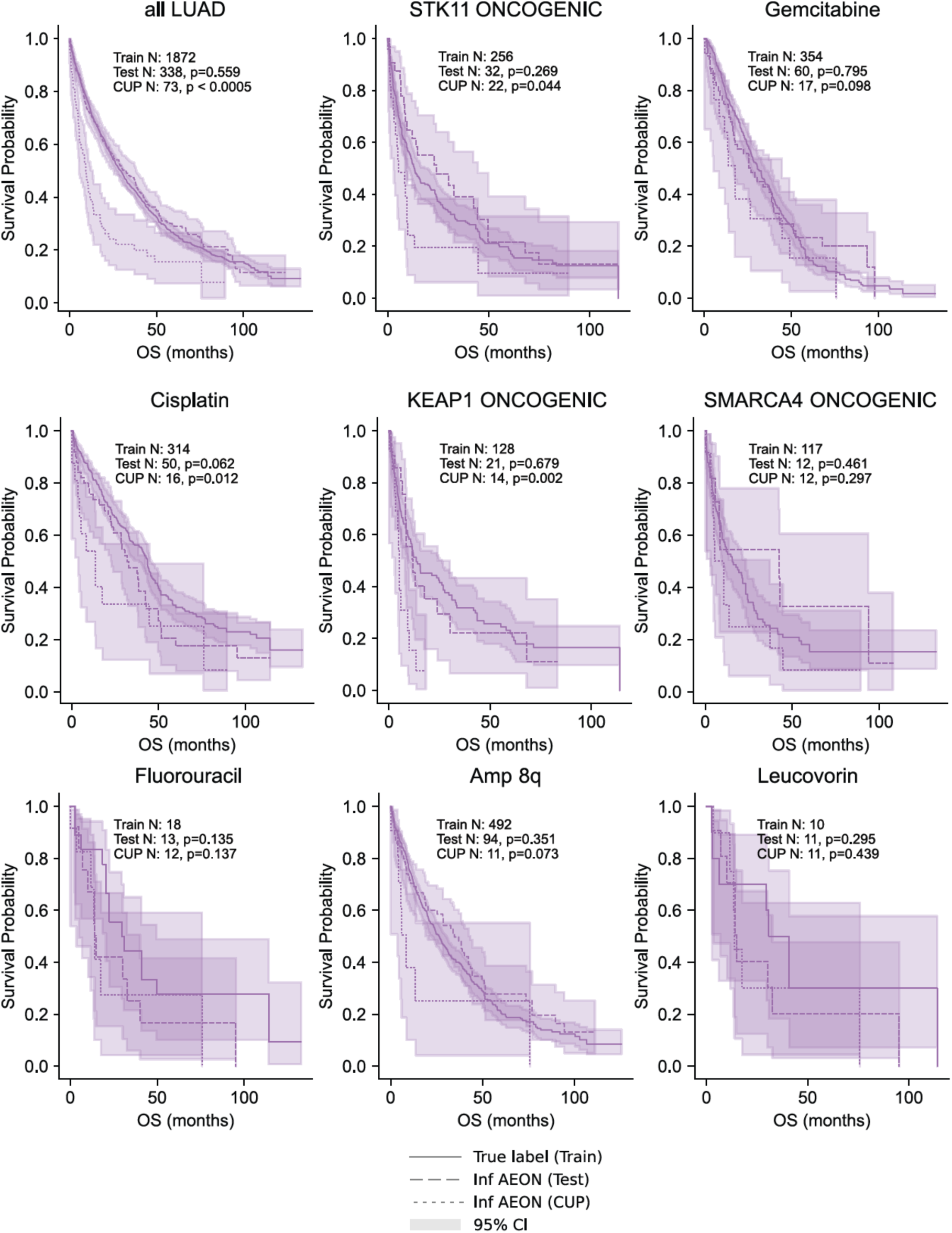
Expanded set of common LUAD alterations and treatment regimes assessed for survival difference correction between LUAD train, test-inferred LUAD and CUP-inferred LUAD subgroups, where at least 10 LUAD-inferred CUP samples are affected by the alteration or treatment indicated.

**Extended Data Fig. 6.**
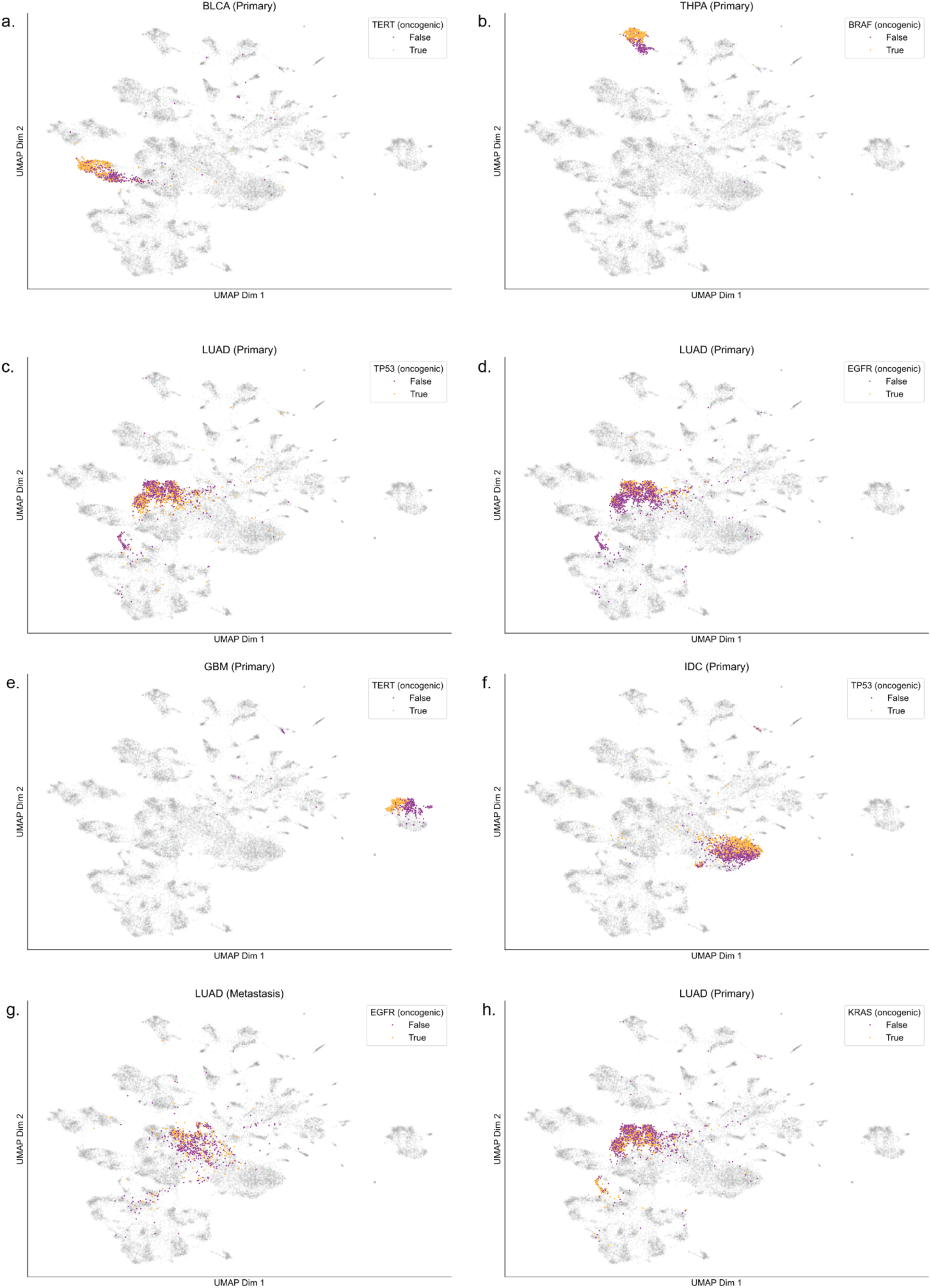
Genomic determinants of histopathologic representation. (a) *TERT* mutational status in primary urothelial carcinoma of the urinary bladder, (b) *BRAF* mutational status in primary papillary thyroid cancer, (c-d) *TP53* and *EGFR* mutational status in primary lung adenocarcinoma, (e) *TERT* mutational status in primary glioblastoma, (f) *TP53* mutational status in primary invasive ductal carcinoma of the breast, (g) *EGFR* mutational status in metastatic lung adenocarcinoma, and (h) *KRAS* mutational status in primary lung adenocarcinoma.

**Extended Data Fig. 7.**
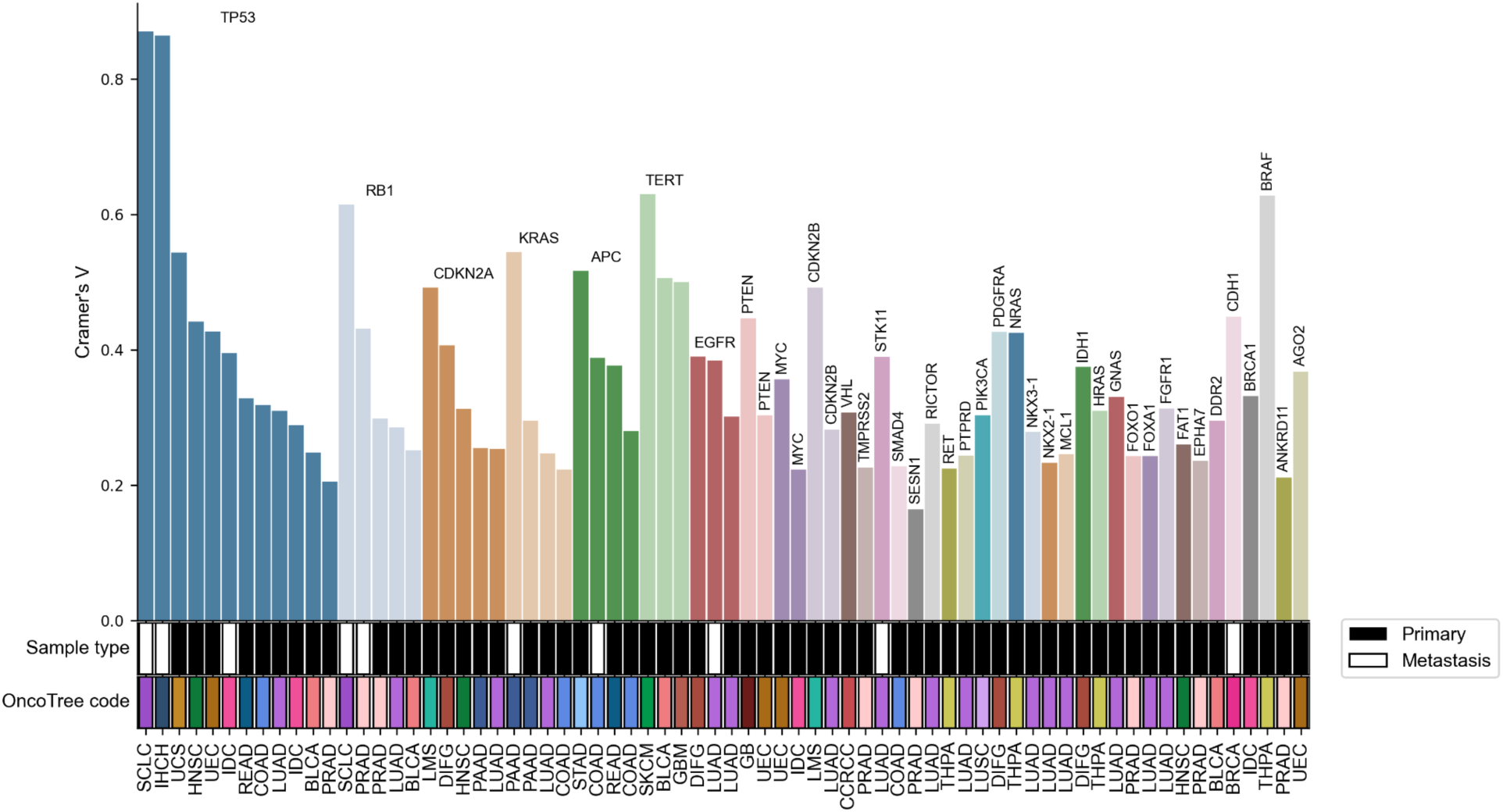
Effect size analysis for oncogenic genomic alterations associated with specific Leiden clusters. All analyses were limited to specific OncoTree codes and sample types. Only Cramer’s V values for significant results by Fisher’s exact test with false discovery rate controlled at 1% are shown.

**Extended Data Fig. 8.**
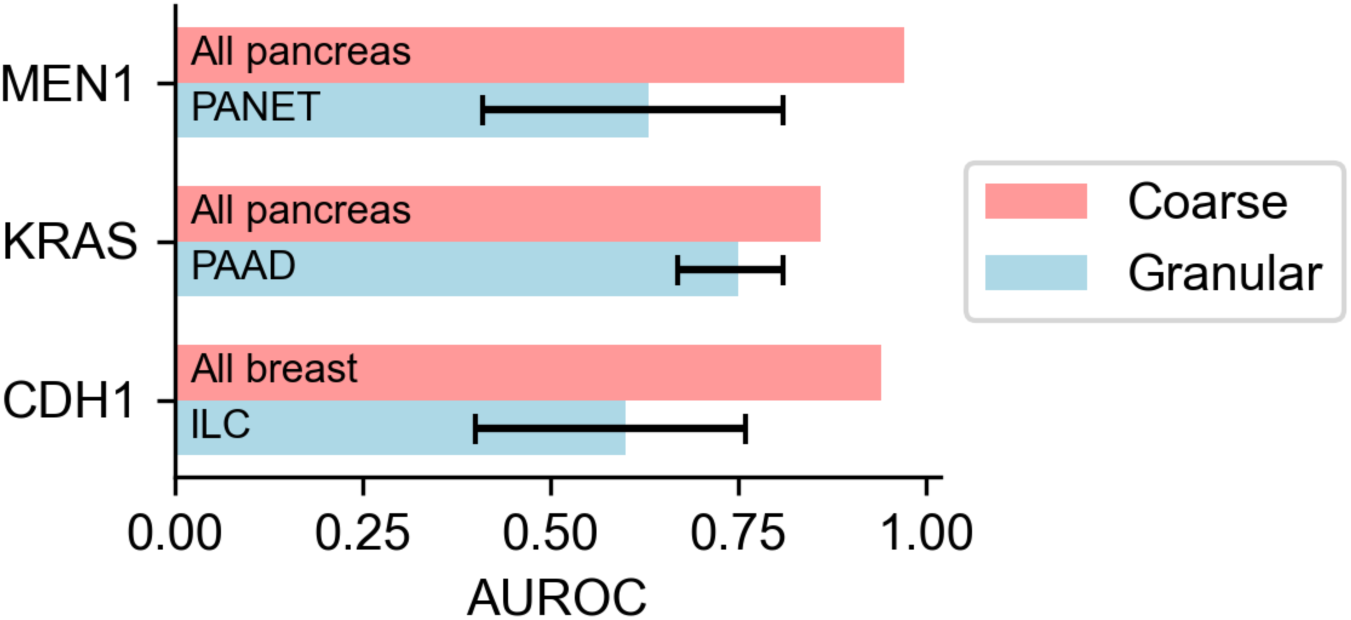
Previously reported phenotypic associations of mutational status are abrogated by limiting to granular cancer subtypes. Error bars denote 95% confidence interval by 1000-fold bootstrapping in the test set.

**Extended Data Figure 9.**
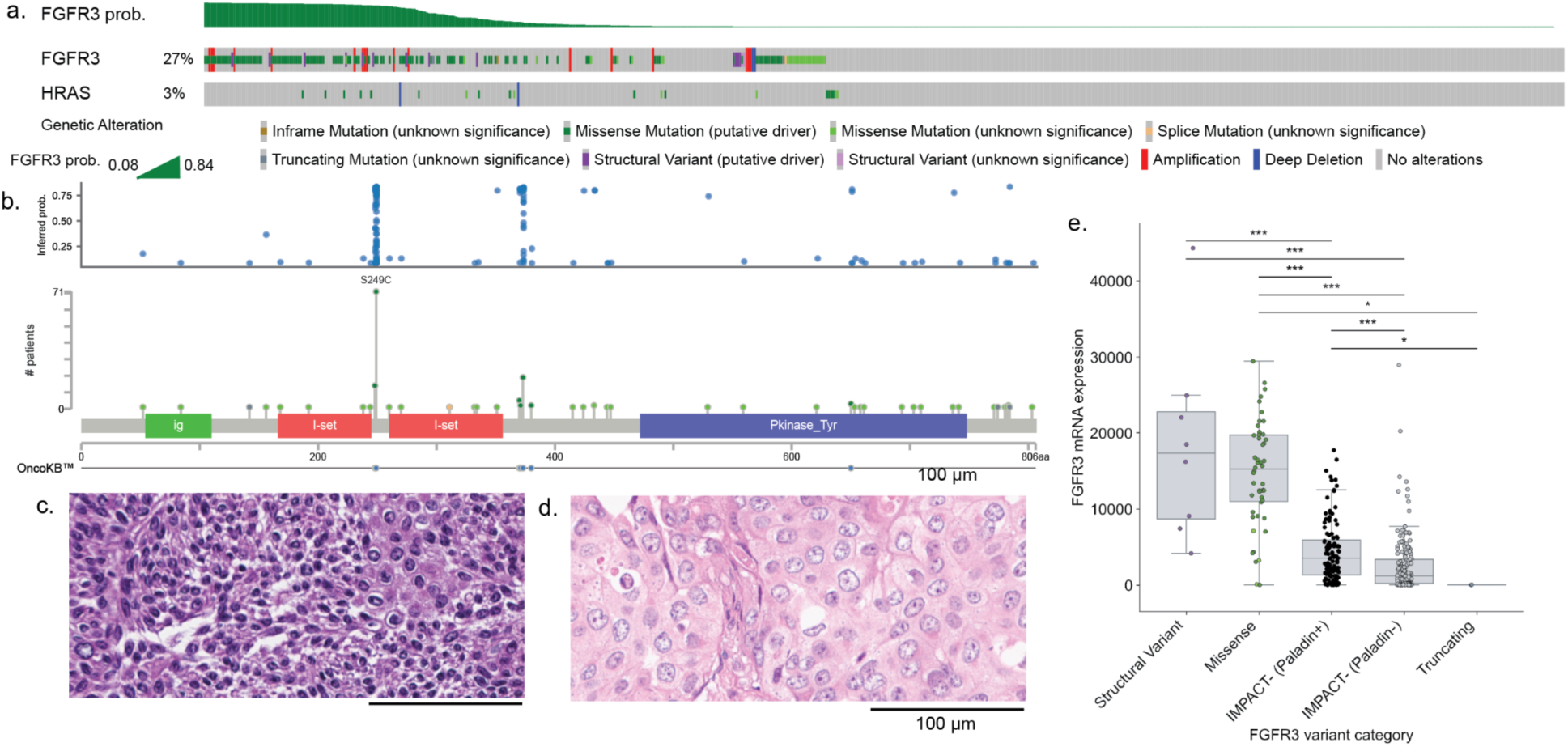
Correlates of inferred alteration probability in *FGFR3*. (a) OncoPrint for *FGFR3*, *FGFR2*, and *HRAS* in urothelial carcinoma. (b) Inferred probability of *FGFR3* oncogenic variant disaggregated by IMPACT-detected variant subtype and locus. (c,d) *FGFR3*-altered (c) and wildtype (d) cases, with altered cases depicting raisinoid nuclei. (e) *FGFR3* transcription versus inferred probability of oncogenic alteration in the external test set.

## Works Cited

1. Kundra, R. et al. OncoTree: A Cancer Classification System for Precision Oncology. JCO Clinical Cancer Informatics (2021) doi:10.1200/CCI.20.00108.

2. Krämer, A. et al. Cancer of unknown primary: ESMO Clinical Practice Guideline for diagnosis, treatment and follow-up. Annals of oncology : official journal of the European Society for Medical Oncology 34, (2023).

3. Varghese, A. M. et al. Clinical and molecular characterization of patients with cancer of unknown primary in the modern era. Annals of oncology : official journal of the European Society for Medical Oncology 28, (2017).

4. Varadhachary, G. R. & Raber, M. N. Cancer of Unknown Primary Site. (2014) doi:10.1056/NEJMra1303917.

5. Raghav, K. Cancer of Unknown Primary Site. New England Journal of Medicine (2025) doi:10.1056/NEJMcp2402691.

6. Darmofal, M. et al. Deep-Learning Model for Tumor-Type Prediction Using Targeted Clinical Genomic Sequencing Data. Cancer discovery 14, (2024).

7. Malone, E. R., Oliva, M., Sabatini, P. J. B., Stockley, T. L. & Siu, L. L. Molecular profiling for precision cancer therapies. Genome Medicine 12, 1–19 (2020).

8. Moon, I. et al. Machine learning for genetics-based classification and treatment response prediction in cancer of unknown primary. Nature medicine 29, (2023).

9. Liu, X. et al. Site-specific therapy guided by a 90-gene expression assay versus empirical chemotherapy in patients with cancer of unknown primary (Fudan CUP-001): a randomised controlled trial. The Lancet. Oncology 25, (2024).

10. Lu, M. Y. et al. AI-based pathology predicts origins for cancers of unknown primary. Nature 594, 106–110 (2021).

11. Wang, X. et al. A pathology foundation model for cancer diagnosis and prognosis prediction. Nature 634, 970–978 (2024).

12. Noorbakhsh, J. et al. Deep learning-based cross-classifications reveal conserved spatial behaviors within tumor histological images. Nature Communications 11, 1–14 (2020).

13. Shao, Z., et al. TransMIL: Transformer based Correlated Multiple Instance Learning for Whole Slide Image Classification. (2021).

14. Li, B., Li, Y. & Eliceiri, K. W. Dual-stream Multiple Instance Learning Network for Whole Slide Image Classification with Self-supervised Contrastive Learning. (2020).

15. Sanchez-Vega, F. et al. Oncogenic Signaling Pathways in The Cancer Genome Atlas. Cell 173, 321–337.e10 (2018).

16. Hoadley, K. A. et al. Cell-of-Origin Patterns Dominate the Molecular Classification of 10,000 Tumors from 33 Types of Cancer. Cell 173, (2018).

17. Liu, J. et al. An Integrated TCGA Pan-Cancer Clinical Data Resource to Drive High-Quality Survival Outcome Analytics. Cell 173, (2018).

18. Alexander, J. et al. Histopathological Identification of Colon Cancer with Microsatellite Instability. The American Journal of Pathology 158, 527 (2001).

19. Kather, J. N. et al. Deep learning can predict microsatellite instability directly from histology in gastrointestinal cancer. Nature Medicine 25, 1054–1056 (2019).

20. Saillard, C. et al. Validation of MSIntuit as an AI-based pre-screening tool for MSI detection from colorectal cancer histology slides. Nature Communications 14, 1–11 (2023).

21. Echle, A. et al. Clinical-Grade Detection of Microsatellite Instability in Colorectal Tumors by Deep Learning. Gastroenterology 159, (2020).

22. Wagner, S. J. et al. Transformer-based biomarker prediction from colorectal cancer histology: A large-scale multicentric study. Cancer cell 41, (2023).

23. Richard Boland, C. & Goel, A. Microsatellite Instability in Colorectal Cancer. Gastroenterology 138, 2073 (2010).

24. Battaglin, F., Naseem, M., Lenz, H. J. & Salem, M. E. Microsatellite instability in colorectal cancer: overview of its clinical significance and novel perspectives. Clinical advances in hematology & oncology : H&O 16, (2018).

25. Liechty, B. et al. Machine learning can aid in prediction of IDH mutation from H&E-stained histology slides in infiltrating gliomas. Scientific Reports 12, 1–12 (2022).

26. Pao, J. J. et al. Predicting EGFR mutational status from pathology images using a real-world dataset. Scientific Reports 13, 1–13 (2023).

27. Arslan, S. et al. A systematic pan-cancer study on deep learning-based prediction of multi-omic biomarkers from routine pathology images. Commun. Med. 4, 48 (2024).

28. El Nahhas, O. S. M., et al. Regression-based Deep-Learning predicts molecular biomarkers from pathology slides. Nat. Commun. 15, 1253 (2024).

29. Kather, J. N. et al. Pan-cancer image-based detection of clinically actionable genetic alterations. Nature Cancer 1, 789–799 (2020).

30. Wang, Y. K., et al. Screen Them All: High-Throughput Pan-Cancer Genetic and Phenotypic Biomarker Screening from H&E Whole Slide Images. (2024).

31. Vang, R., Shih, I.-M. & Kurman, R. J. Ovarian Low-grade and High-grade Serous Carcinoma: Pathogenesis, Clinicopathologic and Molecular Biologic Features, and Diagnostic Problems. Advances in Anatomic Pathology 16, 267 (2009).

32. Pareja, F. et al. A Genomics-Driven Artificial Intelligence-Based Model Classifies Breast Invasive Lobular Carcinoma and Discovers CDH1 Inactivating Mechanisms. Cancer research 84, (2024).

33. Koboldt, D. C. Best practices for variant calling in clinical sequencing. Genome Medicine 12, 1–13 (2020).

34. Solomon, J. P. et al. NTRK fusion detection across multiple assays and 33,997 cases: diagnostic implications and pitfalls. Modern pathology : an official journal of the United States and Canadian Academy of Pathology, Inc 33, 38 (2019).

35. Chakravarty, D. et al. OncoKB: A Precision Oncology Knowledge Base. JCO precision oncology 2017, 10.1200/PO.17.00011 (2017).

36. Suehnholz, S. P. et al. Quantifying the Expanding Landscape of Clinical Actionability for Patients with Cancer. Cancer Discov 14, 49–65 (2024).

37. Boehm, K. M., Khosravi, P., Vanguri, R., Gao, J. & Shah, S. P. Harnessing multimodal data integration to advance precision oncology. Nat. Rev. Cancer 22, 114–126 (2022).

38. Cheng, D. T. et al. Memorial Sloan Kettering-Integrated Mutation Profiling of Actionable Cancer Targets (MSK-IMPACT): A Hybridization Capture-Based Next-Generation Sequencing Clinical Assay for Solid Tumor Molecular Oncology. The Journal of Molecular Diagnostics : JMD 17, 251 (2015).

39. Zehir, A. et al. Mutational landscape of metastatic cancer revealed from prospective clinical sequencing of 10,000 patients. Nature Medicine 23, 703–713 (2017).

40. Pugh, T. J. et al. AACR Project GENIE: 100,000 Cases and Beyond. Cancer Discov 12, 2044–2057 (2022).

41. STK11 and KEAP1 mutations as prognostic biomarkers in an observational real-world lung adenocarcinoma cohort. ESMO Open 5, e000706 (2020).

42. Sanchez-Cespedes, M. et al. Inactivation of LKB1/STK11 Is a Common Event in Adenocarcinomas of the Lung1. Cancer Res 62, 3659–3662 (2002).

43. Guercio, B. J. et al. Clinical and Genomic Landscape of FGFR3-Altered Urothelial Carcinoma and Treatment Outcomes with Erdafitinib: A Real-World Experience. Clinical cancer research : an official journal of the American Association for Cancer Research 29, (2023).

44. Al-Ahmadie, H. A. et al. Somatic mutation of Fibroblast Growth Factor Receptor-3 (FGFR3) defines a distinct morphologic subtype of high-grade urothelial carcinoma. The Journal of pathology 224, 270 (2011).

45. Traag, V. A., Waltman, L. & van Eck, N. J. From Louvain to Leiden: guaranteeing well-connected communities. Scientific Reports 9, 1–12 (2019).

46. Iro, H. & Katabi, N. Salivary Duct Carcinoma. http://dx.doi.org/10.1016/j.path.2016.04.002 doi:10.1016/j.path.2016.04.002.

47. Comprehensive Molecular Portraits of Invasive Lobular Breast Cancer. Cell 163, 506–519 (2015).

48. Rekhtman, N. All That Is Small Is Not a Small-Cell Carcinoma: Thoracic SMARCA4-Deficient Undifferentiated Tumors Masquerading as SCLC. Clinical cancer research : an official journal of the American Association for Cancer Research 30, (2024).

49. Rahadiani, N., Stephanie, M., Perkasa, A. G., Handjari, D. R. & Krisnuhoni, E. p53 expression is associated with tumor stage, grade and subtype in patients with hepatocellular carcinoma. Molecular and Clinical Oncology 19, 54 (2023).

50. Tan, H.-L. et al. Rb Loss is Characteristic of Prostatic Small Cell Neuroendocrine Carcinoma. Clinical cancer research : an official journal of the American Association for Cancer Research 20, 890 (2013).

51. Riley, R. D. et al. Evaluation of clinical prediction models (part 2): how to undertake an external validation study. BMJ 384, (2024).

52. Chen, R. J. et al. Pan-cancer integrative histology-genomic analysis via multimodal deep learning. Cancer cell 40, (2022).

53. Fu, Y. et al. Pan-cancer computational histopathology reveals mutations, tumor composition and prognosis. Nature Cancer 1, 800–810 (2020).

54. Wang, X. et al. AMPK promotes SPOP-mediated NANOG degradation to regulate prostate cancer cell stemness. Developmental cell 48, 345 (2018).

55. Ng, P. K.-S. et al. Systematic Functional Annotation of Somatic Mutations in Cancer. Cancer cell 33, 450 (2018).

56. Gong, M. et al. Loss-of-function mutations in KEAP1 drive lung cancer progression via KEAP1/NRF2 pathway activation. Cell Communication and Signaling 18, 1–11 (2020).

57. Boehm, K. M. et al. Multimodal histopathologic models stratify hormone receptor-positive early breast cancer. Nature Communications 16, 1–14 (2025).

58. Jee, J. et al. Automated real-world data integration improves cancer outcome prediction. Nature 636, 728–736 (2024).

